# Handedness and midsagittal corpus callosum morphology: A systematic meta-analytic evaluation

**DOI:** 10.1101/2021.04.27.441635

**Authors:** René Westerhausen, Marietta Papadatou-Pastou

**Affiliations:** Department of Psychology, University of Oslo, Norway; Biomedical Research Foundation of the Academy of Athens, Athens, Greece; School of Education, National and Kapodistrian University of Athens, Athens, Greece

## Abstract

Following a series of seminal studies in the 1980s, left or mixed hand preference is widely considered to be associated with a larger corpus callosum, influencing the interpretation of findings and various theories related to inter-hemispheric processing, brain lateralisation, and hand preference. Recent reviews of the literature, however, report inconsistencies in the literature and cast doubt on the existence of such an association. The aim of the present study was to systematically evaluate and meta-analytically integrate the available data on the effect of hand preference on corpus-callosum morphology. For this purpose, articles were identified via a search in PubMed and Web of Science databases. Studies reporting findings relating handedness (assessed as hand preference) and corpus-callosum morphology in healthy participants were considered eligible. On the basis of a total of *k* = 25 identified studies, random-effects meta-analyses were conducted considering four different group comparisons found in the literature. That is, studies comparing participants of (a) predominantly right- (dRH) and left-hand preference (dLH), (b) consistent right (cRH) and non-cRH preference, (c) cRH with mixed-hand preference (MH), and (d) cRH with consistent left-hand hand preference (cLH). For none of these meta-analyses did we find a significant effect of hand preference, and narrow confidence intervals suggest that the existence of substantial population effect sizes can be excluded. For example, considering the comparison of dRH and dLH, (summarizing *k* = 14 studies incorporating 1910 dRH and 646 dLH participants) the estimated mean effect size was *g* = 0.016 (95% confidence interval: −0.12 to 0.15). Thus, the common practice of assuming an increase in callosal connectivity based on hand preference is likely invalid.

---

The corpus callosum, formed by a bundle of axons, is essential for the coordination and integration of information processing between and across the two cerebral hemispheres (Gazzaniga, 2000). Consequently, it has long been suggested that inter-individual differences in callosal morphology are related to differences in functional hemispheric asymmetries (Galaburda, Rosen, & Sherman, 1990; Ringo, Doty, Demeter, & Simard, 1994; Witelson & Nowakowski, 1991). Recent neuroimaging studies in general support this notion as they have shown an association between measures of structural corpus callosum connectivity and the distribution of neuronal processing between the hemispheres (e.g., Friedrich et al., 2020; Haberling, Badzakova-Trajkov, & Corballis, 2011; Josse, Seghier, Kherif, & Price, 2008; Karolis, Corbetta, & De Schotten, 2019; Labache et al., 2020; Moffat, Hampson, & Lee, 1998; Westerhausen et al., 2006).

Handedness, arguably the most salient functional asymmetry with almost 9 out of 10 individuals being right-handed (Papadatou-Pastou et al., 2020), has also been frequently related to differences in corpus callosum structure and function (for review see, Beaton, 1997; Budisavljevic, Castiello, & Begliomini, 2020). The origin for this line of research can be found in a series of seminal publications by Sandra Witelson (Witelson, 1985, 1989; Witelson & Goldsmith, 1991). She had measured the midsagittal corpus callosum in brain specimen of deceased cancer patients of known hand preference and found the corpus callosum to be larger in mixed-handed individuals than in consistent right-handers. Interpreting callosal size as an indicator of the strength of hemispheric connectivity, these findings were taken to indicate that a less-lateralized hemispheric organization for hand preference was associated with stronger or more efficient connectivity than a strongly lateralized organization. Consequently, studies frequently refer to Witelson’s findings when explaining behavioral or cognitive differences between right- and non-right handers that might be linked to the corpus callosum, even without measuring the corpus callosum itself (some recent examples, see e.g., Parker, Parkin, & Dagnall, 2017; Roberts, Fernandes, & MacLeod, 2020; Sala, Signorelli, Barsuola, Bolognese, & Gobet, 2017; Zapała et al., 2020). For example, callosal size differences have been used to explain superior episodic-memory performance in mixed-as compared to consistent (right)-handers (Prichard, Propper, & Christman, 2013). However, Witelson’s original findings are not necessarily supported by more recent studies and reviews typically summarize the literature as being inconsistent, questioning the existence of a general association (Beaton, 1997; Budisavljevic et al., 2020).

Thus, the aim of the present study was to revisit the question of an association of handedness and corpus callosum morphology using meta-analytic methods. For this purpose, we identified all publications comparing midsagittal corpus callosum morphology in neurological healthy participants based on their hand preference. That is, studies were included if they assessed hand preference by self-report (i.e., questionnaire) and used measures of total or subsectional corpus callosum morphology (as volume, area, or thickness) as a dependent variable. As the studies varied in their definition of handedness groups, we could not integrate all data into a single meta-analytic comparison. Rather, we conducted meta-analyses for four different group comparisons: Studies that compared participants of (a) dominantly right- (dRH) and dominantly left-hand preference (dLH), (b) consistent right (cRH) with non-cRH preference (NcRH), (c) cRH with mixed-hand preference (MH), and (d) cRH with consistent left hand preference (cLH). Thus, meta-analyses (a) and (d) focus on the effect that the direction of hand preference has on the corpus callosum, while meta-analyses (b) and (c) also consider consistency of hand preference. Furthermore, we investigated the presence of heterogeneity and small study bias in this literature. Finally, as the interaction of sex and hand preference has received substantial attention in the literature comparing cRH and NcRH samples when explaining total and isthmal corpus-callosum size (Clarke & Zaidel, 1994; Habib et al., 1991; Witelson, 1989), we conducted a moderator analysis of sex for these comparisons in addition to the overall meta-analysis.

## Material and methods

### Study identification

The study selection was based on a literature search that was conducted on 01. February 2021 on PubMed and Web of Science (Core Collection). The search in PubMed was performed with the search query (“corpus callosum”[All Fields]) AND (“handedness”[All Fields] OR “hand preference” [All Fields]). The search in Web of Science used the search terms (ALL FIELDS: (Corpus callosum) AND ALL FIELDS: (handedness OR hand preference)). Additional studies were identified from the reference list of the selected empirical articles and previous review articles (Beaton, 1997; Budisavljevic et al., 2020; Driesen & Raz, 1995) and by contacting authors who recently had published corpus callosum data on handedness.

### Record screening, article eligibility check, and inclusion

After removing duplicates, potential relevance of the study was step-wise evaluated as described in the following. Firstly, title and abstracts were screened to check whether morphological corpus-callosum measures were utilized and that the sample consisted of human participants. If this was the case, the full-text articles were inspected to verify that corpus callosum measures were reported for non-right handed individuals as well. This step led to the exclusion of studies which had used handedness as an exclusion criterion (typically, excluding non-right handers) or to match participants (without reporting the means for handedness groups). In case of clinical studies, only the data of healthy control groups were considered relevant. In a final evaluation round, it was determined whether sufficient information was available to include the data in the quantitative meta-analysis. If this was not the case the corresponding authors were contacted to obtain the relevant data where possible.

The following list provides an overview of all criteria for screening and eligibility assessment:

- Study languages: publications in English, German, and French were considered
- Species: only data from human (*homo sapiens*) samples was included
- Health: only data from neurologically healthy individuals was considered. Of note, Witelson’s seminal work examined patients suffering from peripheral (i.e., not directly affecting the central nervous system) cancer (Witelson, 1985, 1989; Witelson & Goldsmith, 1991) and was included in the analysis.
- Hand preference had to be assessed using self-report measures, i.e. questionnaires or self-identification. Studies using performance or skill measures for assessment were not included due to sparsity.
- Corpus callosum had to be assessed morphologically regarding its midsagittal shape. Studies reporting raw thickness, area, and volume measures were included.
- Reporting of useable arithmetic data (means and standard deviation or standard error) per group or test statistics (e.g, a t-value) for the group comparison.

An overview of the selection procedure and the number of studies excluded on each step can be found in flow-chart (Fig. 1). The data screening was done by one reviewer.

**Fig. 1.**
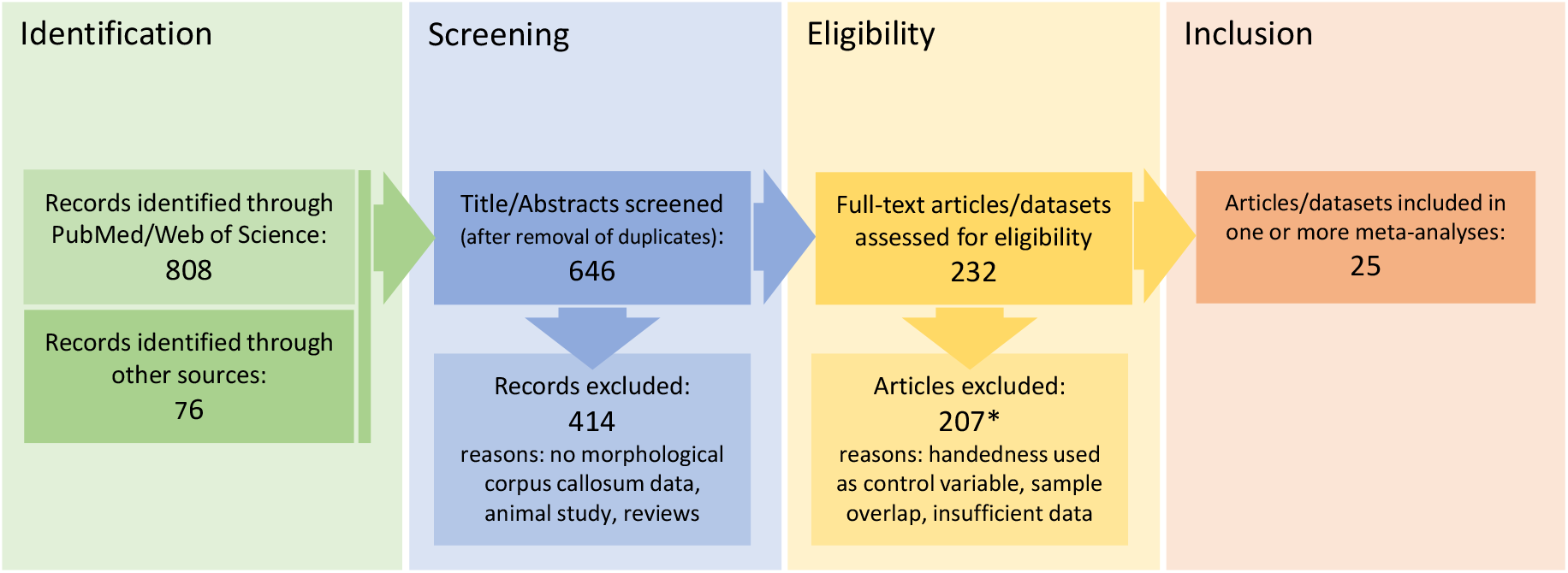
Overview of study identification, screening, and eligibility assessment. All included studies and datasets are presented in Table 1. All studies that were initially considered eligible but did not provide sufficient information for a statistical inclusion or were redundant (i.e., sample overlap) to other included reports can be found in Supplement Table S5 (with reason for the exclusion).

### Data extraction

In order to assure independence of observation, the overlap in samples across studies was evaluated and it was made sure that each sample was included only once per meta-analysis. In case of such overlap, only most complete data (e.g., the largest sample) was included. For example, the data reported by Cowell and Gurd (2018) represents a sample extension of the data published earlier (Gurd et al., 2013) thus only data from the larger 2018 sample was included. A second example, Witelson and Goldsmith (1991), reports the results only of an extension of the male sample compared to the Witelson (1989) article, therefore the male data was taken from the later and the female data from the earlier publication.

Regarding hand preference, the method of assessment and potential thresholds for group formation were extracted together with the number of individuals in each group. Of note, as is typical in the handedness literature (Ocklenburg & Güntürkün, 2018), the studies included used various criteria to define groups of hand preference. Four main approaches were common in the available literature and accordingly considered in the present set of meta-analyses.

- A classification based on the “handedness direction” so that the overall preference across the activities assessed by a handedness questionnaire determines the classification. Typically, a laterality quotient (LQ) of 0 (equivalent to no hand preference, see e.g., (Oldfield, 1971)) was used to split the sample into a group of dRH and dLH individuals.
- Consistency of hand preference across tasks was considered, by comparing cRH individuals with all other (referred to as NcRH). For this purpose, consistency was defined by the primary studies either by arbitrarily setting a high LQ (e.g., LQ > 80% in (Habib et al., 1991)) as cut-off or by a qualitative analyses of the response pattern in the handedness questionnaire. The latter approach follows the suggestion by Witelson (1989), who classified participants as cRH if the answers in Annett’s questionnaire (Annett, 1970) were “all ‘right’, or ‘right’ with some ‘either’ preferences”. All other individuals (i.e., LQ < 80 %; or indicating any ‘left’ preferences) were consequently defined as not being consistent (NcRH).
- Witelson originally picked this solution, as she was not able to find a sufficiently large sample of consistent left-handers to form their own group (Witelson, 1985). However, later researchers did so, resulting in the possibility of comparing a cRH group with a group of MH and cLH individuals, as third and fourth comparison, respectively. That is, the NcRH group as defined above, was split into two groups, whereby typically a high negative LQ threshold (e.g., LQ < −80%) was utilized to separate cLH from the MH group.

Considering the corpus callosum, raw mean and standard deviation of the total corpus callosum were extracted from the primary articles. The focus on raw measures was necessary as a correction for brain size measures was rarely reported, preventing us from conducting a separate analysis considering brain-size differences. For each study, the method of obtaining the data (i.e., post mortem study or in-vivo MRI; including field strength), the method of callosal segmentation, and the measurement type of the dependent variable (i.e., thickness, area, or volume) was recorded. Where available also measures of corpus callosum subsections were extracted. As the method of dividing the corpus callosum into subsections varies between studies, an attempt was made to apply a common frame of reference across studies. That is, we used the geometrical subdivision schema suggested by Jäncke et al. (Jancke, Staiger, Schlaug, Huang, & Steinmetz, 1997), which divides the midsagittal corpus callosum surface into four subsections based on the anterior-posterior extend of the structure (see Fig. 2). This schema was considered optimal, as it is sufficiently broad to subsume some of the more fine-grained alternative subdivision schemas, but also includes an isthmus section, which has received particular attention by the literature following Witelson’s original findings (see e.g., Denenberg, Kertesz, & Cowell, 1991). A transfer was made by applying the Jäncke et al. (1997) subdivision scheme on the visual representation of the callosal subdivision provided in the literature and determine correspondence between subdivisions. An illustration of the procedure and the transfer heuristics used can be found in Supplemental material, Fig. S1.

**Fig. 2.**
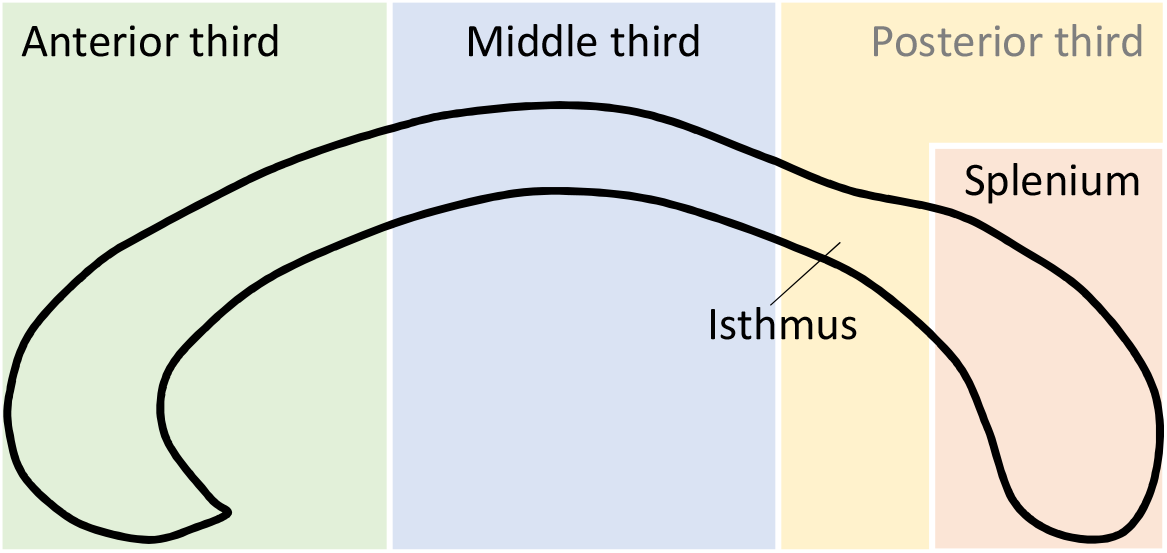
Illustration of corpus callosum subdivision used in the meta-analysis. The approach followed the straight-line method introduced by Jancke et al. (1997). The outline of the corpus callosum (black line) is divided into thirds relative to its anterior-posterior extend. The posterior third is additionally split into a posterior fifth (i.e., the splenium) and the isthmus.

The required corpus callosum data for the meta-analysis was not always readily available in the articles, whereby the following cases were encountered. Firstly, the article did report mean and dispersion of the callosal measures by subgroups (e.g., for female righthanders and male right-handers) but not the entire handedness group. In this case, the data was pooled across the subsamples considering the subsample sizes as weights. Studies where this was the case are indicated in Supplement Tables S1 to S4. Secondly, the article did not report the relevant data, but the author contact was successful. Here the relevant data was either calculated based on the received raw data or the authors were so kind to provide the relevant means and standard deviations. For these cases, a detailed information of the data obtained, the conducted calculations, and results are provided as Supplementary Material Section 3. Thirdly, studies did not provide relevant data and authors were non-responsive or no longer traceable. Here, the study was not included in the meta-analysis. A complete list of these studies can be found in Supplement Table S6.

During data extraction, a within study risk-of-bias assessment was done by focussing on the criteria of selective reporting of results. Specifically, it was checked whether the potentially available corpus-callosum data was also reported (see Supplement Table S5). For example, several studies reported to assess multiple corpus-callosum subsections but only report the findings for the significant comparisons (e.g., Hines, Chiu, McAdams, Bentler, & Lipcamon, 1992; Moffat et al., 1998). If this was the case, the information was noted and considered when evaluating the outcome of the meta-analyses.

Finally, the sex distribution and age range of each study’s sample were recorded in addition to variables describing the assessment of handedness and corpus callosum morphology. An overview of these variables can be found in Table 1 as well as in Supplement Tables S1 to S4. The data extraction was done by one rater.

**Table 1.**
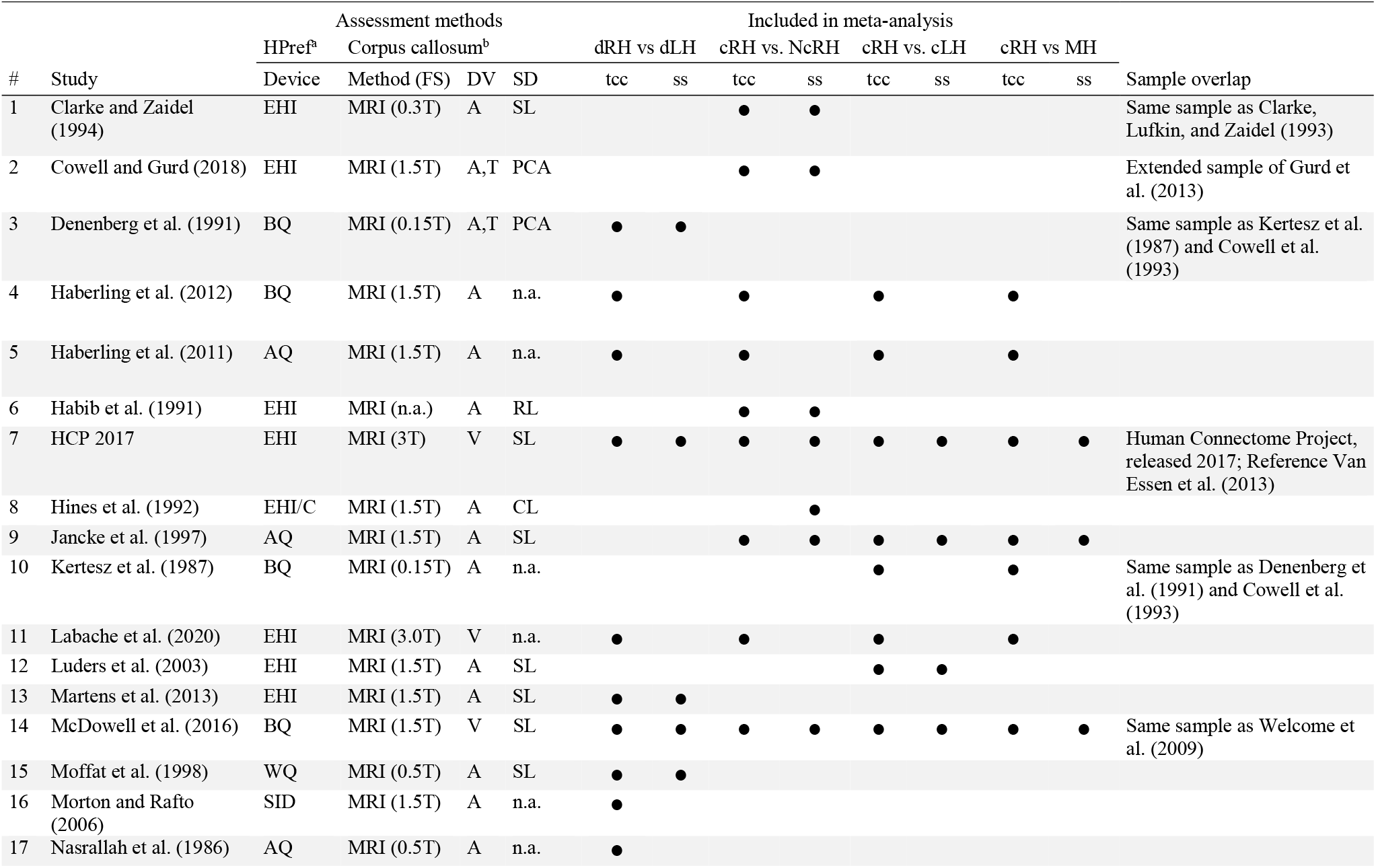
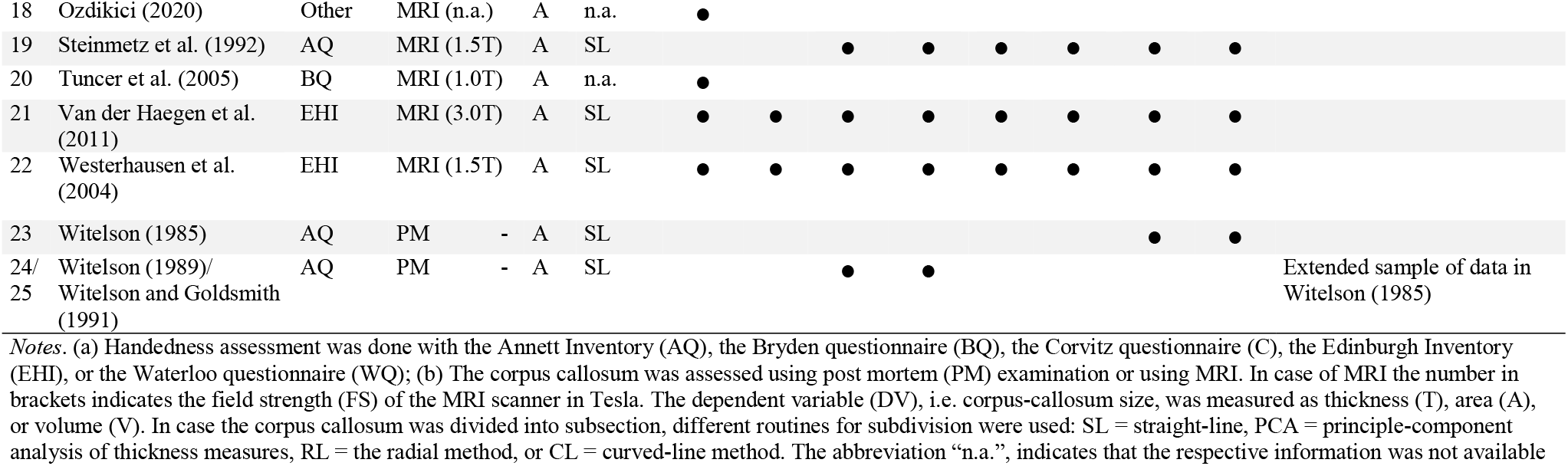
Overview of study inclusion (•) in the meta-analyses considering the total corpus callosum (tcc) or callosal subsections (ss)

### Statistical analysis

The above-outlined study differences in the handedness classification could not allow for an overall analysis without losing relevant information. We therefore decided to conduct a set of four meta-analyses for the following group comparisons: (a) dRH vs. dLH, (b) cRH vs. NcRH, (c) cRH vs. cLH, and (d) cRH vs. MH. Considering (b) we additionally conducted a moderator analysis to explore a suggested interaction of sex and hand preference. The effect size measure Hedges’ *g*, reflecting the standardised mean difference of corpus callosum size, was calculated for each study. In all analyses, the pooled effect size was determined assuming a random-effects model. A random-effects model was preferred, as a fixed-effect model was considered inappropriate given the variability in callosal and handedness measures between studies. The between study variance (*τ*^2^) was estimated using the DerSimonian-Laird approach and the Hartung-Knapp adjustment was chosen to account for the small number of studies included.

As the number of studies analysing callosal subsections was smaller than the number of studies included in the analysis of the total corpus callosum, we used power analysis to determine the minimal number of studies (*k_min_*) required to provide a power of at least 0.80 for a population effect size of *d* = 0.2 (i.e., a small effect according to Cohen, 1992). For this purpose, we used the average sample size per study and the heterogeneity estimates from the total corpus callosum meta-analyses as best guess values for the subsection analyses. The number of required studies was then determined by iteratively applying the *power. analysis* script provided with the “dmetar” R package (version 0.0.9; (Harrer, Cuijpers, Furukawa, & Ebert, 2019) with the above parameters. From this it was determined that the minimum number of studies for comparison (a) was *k_min_* = 8, for (b) *k_min_* = 8, for (c) *k_min_* = 12, and for (d) *k_min_* = 7. A meta-analytic integration was only performed when the available number of studies was at least equal to the determined *k_min_*.

Visual examination and Egger’s regression were used to inspect the funnel plots of each meta-analysis for a potential small study bias. Also, if one study had a weight in the analysis of 25% or above, the meta-analysis was repeated without this study to evaluate the stability of the population-effect size, as a form of sensitivity analysis.

All effect size calculations were done using the “esc” R package (version 0.5.1; (Lüdecke, 2018). Meta-analytic procedures were conducted using functions of the libraries “metafor” (version 2.4; (Viechtbauer, 2020), “meta” (version 4.16-1; (Schwarzer, 2020), and “dmetar” (version 0.0.9; (Harrer et al., 2019) using the R environment (version 4.0.3). The reporting of the meta-analysis followed the PRISMA check list (Page et al., 2021), but was not pre-registered.

### Data availability

The extracted mean values and standard deviations that served as the basis of the present meta-analyses are available in tabulated form on the accompanying OSF platform (https://osf.io/sw6ev/). All R scripts used for running the presented analyses are also available there, and allow for reproduction of all results. The PRISMA checklist is also available on this platform. The meta-analysis has not been preregistered.

## Results

### Descriptive statistics of included studies

Data from *k* = 25 reports and datasets published between 1985 and 2020 were included in one or several meta-analyses as indicated in Table 1. In 10 (40%), 7 (28%), and 5 (20%) of these studies, hand preference was assessed with versions of the Edinburgh Handedness Inventory (Oldfield, 1971), the Annett questionnaire (Annett, 1970), or the handedness questionnaires suggested by Bryden (1977), respectively. A total of 24 (96%) studies provided measures of total corpus callosum size and 17 (68%) studies provided measures for one or more callosal subsections (of note, one study only provided subsection data; see Table 1 for details). The corpus callosum was assessed by post-mortem morphometry in 3 (12%) of these reports and using in-vivo magnetic-resonance imaging in 22 (88%) studies. Callosal size was measured as midsagittal area in 22 (88%) studies and as volume in 3 (12%) studies. Thickness measures were additionally reported in 2 (8%) of the included studies.

### Meta-analysis set for the dRH vs. dLH comparison

Concerning the total corpus callosum area, effect sizes from *k* = 14 studies were included in the meta-analysis, which incorporated data from 1910 dRH and 646 dLH participants. Details on the included studies and data extraction can be found the Supplement Table S1. As shown in the forest plot (Fig. 3), the estimated mean effect size *g* = 0.016 did not deviate significantly from zero (*t* = 0.27, *p* = 0.79). The 95% confidence interval (CI95%) for *g* ranged from −0.12 to 0.15. The between study variance was *τ*^2^ = 0.01, suggesting a 95% prediction interval from −0.24 to 0.27 around the mean effect. Neither inspection of the funnel plots (see Supplement Fig. S2) nor the Egger’s test of the intercept (*a* = −1.35, *t* = −2.01, *p* = 0.07) suggested a small study bias.

**Fig. 3.**
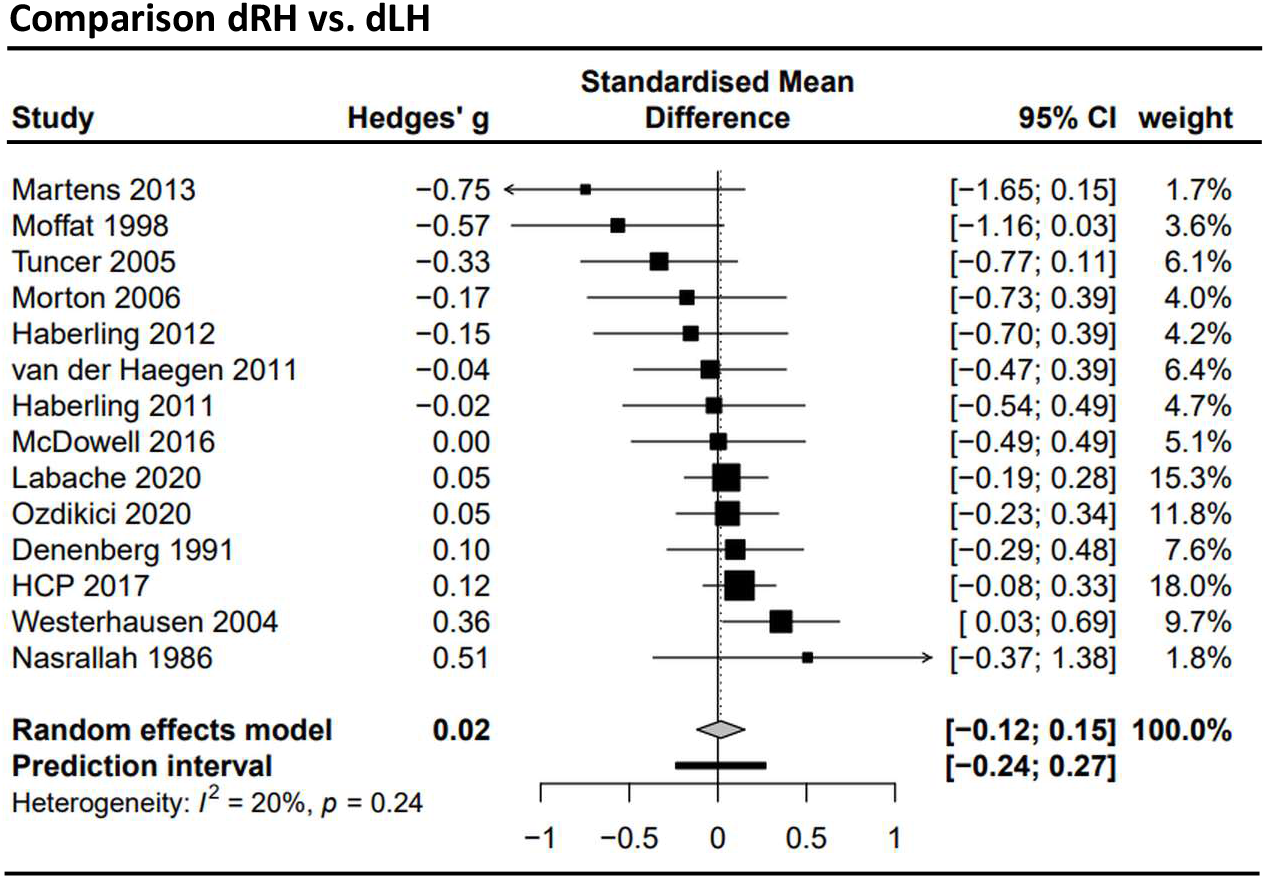
Forest plot of the meta-analysis of studies comparing dominant right (dRH) and dominant left (dLH) samples (dependent variable: total corpus callosum size). The total sample size across all *k* = 14 studies was *n* = 1910 for dRH and *n* = 646 for dLH sample. Negative values indicate the dLH group to have a larger corpus callosum, positive values indicate the dRH group to have a larger corpus callosum.

Meta-analyses of the subsectional data were not conducted as the number of studies available was below the minimum determined by the a priori power analysis. An overview of the available studies and their effect sizes can be found in Supplement Fig. S4.

### Meta-analysis set for the cRH vs. NcRH comparison

Data from *k* = 13 studies (see Supplement Table S2) with a total of 1175 cRH and 1148 NcRH participants were included in the meta-analysis regarding the total corpus callosum area. The estimated mean effect size *g* = −0.077 (CI95%: −0.26; 0.11) did not deviate significantly from zero (*t* = −0.89, *p* = 0.39; see forest plot Fig. 4). The heterogeneity between studies was *τ*^2^ = 0.03 resulting in a prediction interval from −0.52 to 0.37. Inspection of the funnel plots (see Supplement Fig. S2) and the Egger’s test (*a* = −1.02, *t* = −1.41, *p* = 0.19) did not indicate a small study bias.

**Fig. 4.**
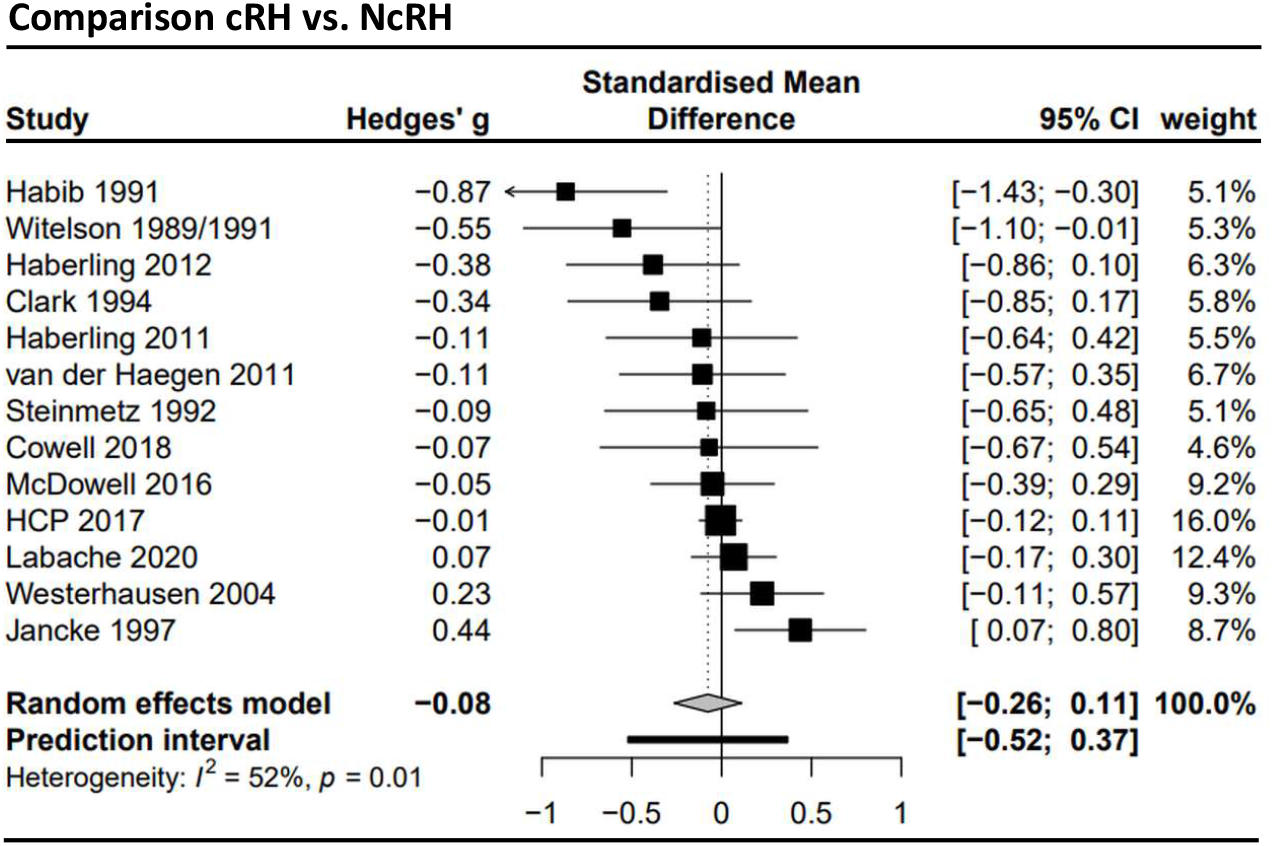
Forest plot of the meta-analysis of studies comparing consistent right-handers (cRH) and non-cRH (NcRH) samples (dependent variable: total corpus callosum size). The analysis included *k* = 13 studies with a total sample of *n* = 1175 for dRH and *n* = 1148 for NcRH. Negative values indicate the NcRH group to have a larger corpus callosum, positive values indicate the cRH group to have a larger corpus callosum.

Meta-analyses for all four subsections were conducted, but none suggested a significant group effect (see Fig. 5). Concerning the anterior third subregion, the metaanalysis included *k* = 9 studies (cRH: *n* = 972; NcRH: *n* = 875) and estimated a mean effect of *g* = −0.086 (CI95%: −0.32 to 0.15; *t* = −0.84, *p* = 0.42; *τ*^2^ = 0.03; prediction interval: −0.57 to 0.40). For the middle third, the analysis included *k* = 10 studies (cRH: *n* = 991; NcRH: *n* = 908) and estimated a mean effect of *g* = 0.03 (CI95%: −0.15 to 0.22; *t* = 0.41, *p* = 0.69; *τ*^2^ = 0.02; prediction interval: −0.36 to 0.42). As in this analysis the HCP sample dominate the overall analysis (weight of 25%), we recalculated the analysis without the HCP sample. Here the mean effect size was estimated to be *g* = 0.05 (CI95%: −0.18 to 0.27; *t* = 0.48, *p* = 0.64; *τ*^2^ = 0.04). Regarding the isthmus, *k* = 11 studies (cRH: *n* = 1011; NcRH: *n* = 923) yielded a mean effect of *g* = −0.032 (CI95%: −0.21 to 0.15; *t* = −0.39, *p* = 0.71; *τ*^2^ = 0.02; prediction interval: −0.40 to 0.34). Finally, the mean effect for the splenium region (*k* = 10; cRH: *n* = 991; NcRH: *n* = 908) was *g* = −0.049 (CI95%: −0.22 to 0.12; *t* = −0.66, *p* = 0.53; *τ*^2^ = 0.01; prediction interval: −0.36 to 0.26). As the HCP sample with a 31.4% weight dominated the analysis, we also estimated the mean effect excluding the HCP sample: *g* = −0.08 (CI95%: −0.30 to 0.14; *t* = −0.85, *p* = 0.42; *τ*^2^ = 0.03).

**Fig. 5.**
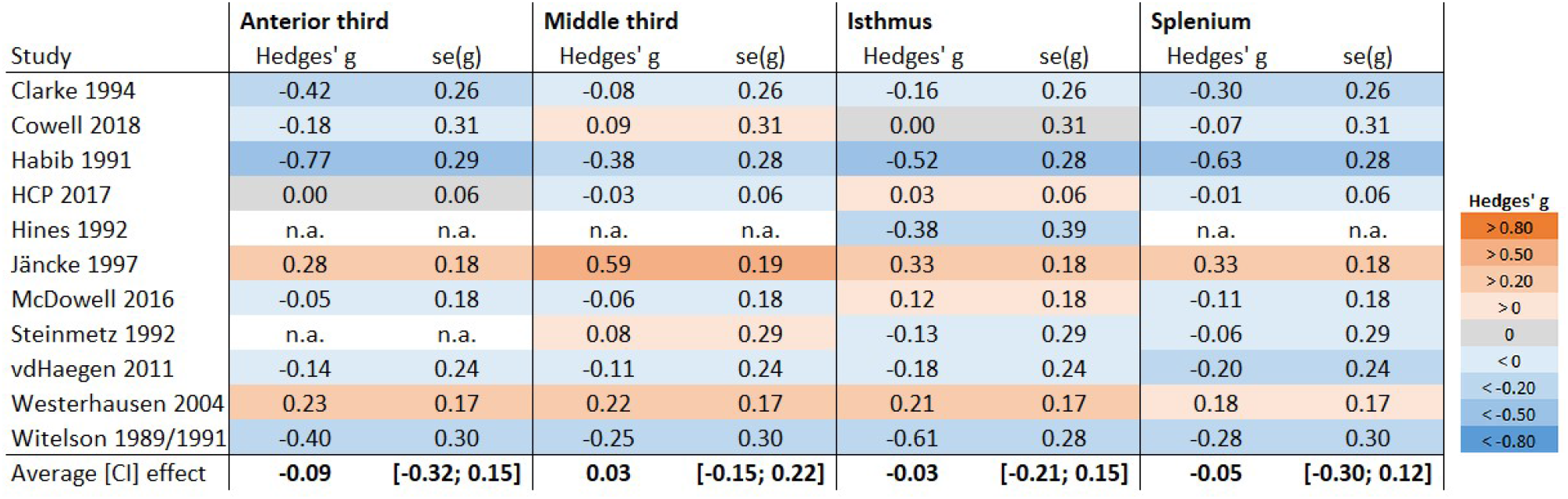
Overview of subsection meta-analyses comparing cRH and NcRH samples. The graph presents the effect size (Hedges’ g) and standard error of the effect size (*se(g)*) for each study. Negative values indicate the subsection to be larger in the NcRH group, positive values indicate the subsection to be larger in the the cRH group. The provided average effect is estimated using a random-effects model. The values in brackets are the 95% confidence interval. Color coding was based on the Cohen’s effect-size heuristics (Cohen, 1992) as indicated in the figure legend. Note, for some studies data was not available (n.a.) for some of the subsections.

The additional conducted subgroup analyses did neither for the total corpus callosum (*Q* = 1.02; *df* = 1; *p* = 0.31) nor for the isthmus subsection find a significant difference between the groups (*Q* = 0.82; *df* = 1; *p* = 0.37). Regarding the total corpus callosum, the analysis included *k* = 11 datasets of a female subgroup, which was characterized by a *g* = 0.06 (CI95%: −0.04 to 0.15; *τ*^2^ < 0.001). The male subgroup analysis contained *k* = 10 datasets, yielding a *g* = −0.11 (CI95%: −0.46 to 0.25) but showing a comparatively high between study variance (*τ*^2^ = 0.09). Regarding isthmus, the female analysis included *k* = 11 datasets and suggested a mean effect size of *g* = 0.06 (CI95%: −0.11 to 0.23; *τ*^2^ = 0.01). The male subgroup analysis (*k* = 9 datasets) found a *g* = −0.10 (CI95%: −0.48 to 0.27; *τ*^2^ = 0.09). The forest plot of both analyses are presented in Supplement Fig. S3.

### Meta-analysis set for the cRH vs. cLH comparison

The data of *k* = 11 studies (see Supplement Table S3) was included in the cRH vs. cLH metaanalysis summarizing the data from a total of 1142 cRH and 306 cLH participants. The group difference in total corpus callosum size was estimated to be *g* = 0.06 (CI95%: −0.10; 0.23) and did not deviate significantly from zero (*t* = 0.87, *p* = 0.41). As can be seen in the forest plot (Fig. 6), the heterogeneity between studies was with *τ*^2^ < 0.01 comparatively low leading to a narrow prediction interval of −0.12 to 0.25. Neither funnel plots nor Egger’s regression (*a* = −1.08, *t* = −1.01, *p* = 0.34) suggested a small study bias. Subsectional meta-analyses were not conducted as the number of studies available was below the minimum determined by the power analysis (see Supplement Fig. S4 for an overview).

**Fig. 6.**
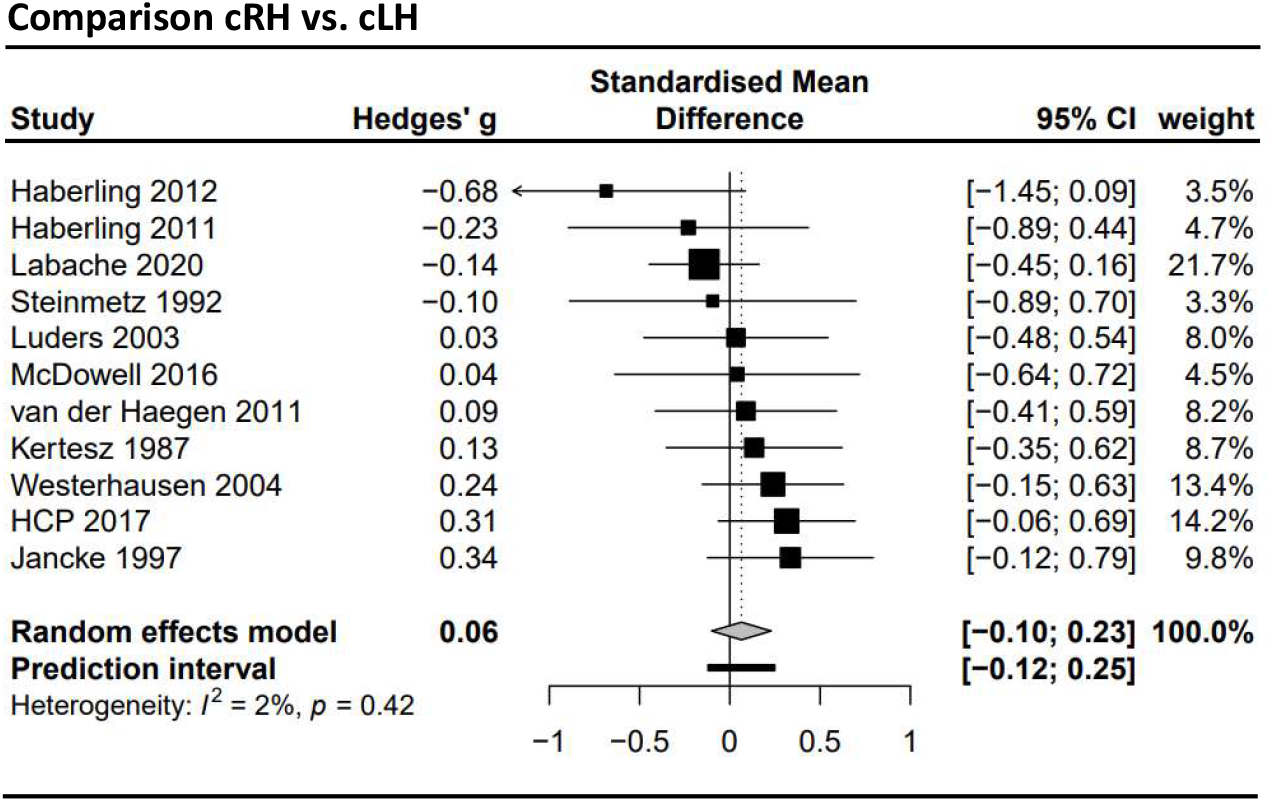
Forest plot of the meta-analysis of studies comparing consistent right-handers (cRH) and consistent left-handers (cLH) samples (dependent variable: total corpus callosum size). It included *k* = 11 studies with a total sample of *n* = 1142 cRH and *n* = 306 cLH participants. Negative values indicate the cLH group to have a larger corpus callosum, positive values indicate the cRH group to have a larger corpus callosum.

### Meta-analysis set for the cRH vs. MH comparison

The meta-analysis contrasting cRH and MH groups integrated data from *k* = 11 studies (see Supplement Table S4 and forest plot in Fig. 7) with data from a total of 1139 cRH and 810 MH participants. The differences in total corpus callosum size between the groups was estimated to *g* = −0.004 (CI95%: −0.20; 0.19) and was not statistically significant (*t* = −0.05, *p* = 0.96). Between study variance was estimated to *τ*^2^ = 0.02 so that the 95% prediction interval was −0.40 to 0.40. No indication for a small study bias was found (Egger’s regression: *a* = −0.06, *t* = −0.08, *p* = 0.94; see funnel plot in Supplement Fig. S2). A subsection analysis was only conducted for the splenium subsection (*k* = 7; cRH: *n* = 909; MH: *n* = 661). The mean effect size for the splenium analysis was *g* = 0.0005 (CI95%: −0.28 to 0.28; see also Supplement Fig. S4) and was not significant (*t* = 0.00, *p* = 0.996; *τ*^2^ = 0.03; prediction interval: −0.55 to 0.56).

**Fig. 7.**
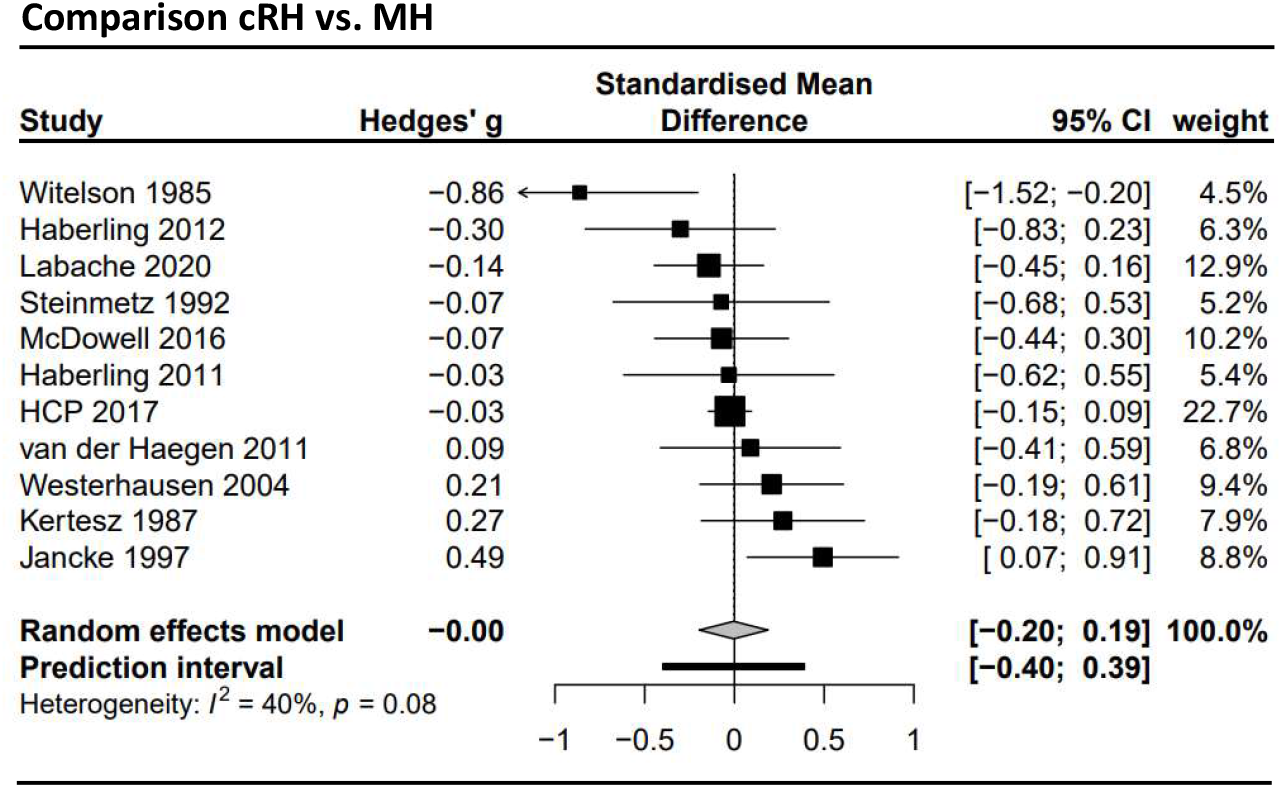
Forest plot of the meta-analysis of studies comparing consistent right-handers (cRH) and mixed-handers (MH) samples (dependent variable: total corpus callosum size). The analysis included *k* = 11 studies with a total sample of *n* = 1139 cRH and *n* = 810 MH participants. Negative values indicate the MH group to have a larger corpus callosum, positive values indicate the cRH group to have a larger corpus callosum.

## Discussion

The present study investigated the presence of differences in corpus-callosum morphology between individuals of different hand preference, using meta-analytic techniques. Irrespective of the nature of the compared hand preference groups, we did not find a significant association of handedness and corpus-callosum morphology, neither for total corpus callosum size, nor for any of its subsections.

The interpretation of these non-significant findings requires first to consider the sensitivity of the conducted analyses. Considering the confidence limits, population effects (|*g*| > 0.2) in total corpus callosum size can be excluded both for the comparison of dRH with dLH and the comparison of cRH with MH. Thus, effects explaining more than 1% variance in the data can thus be excluded for those analyses. For the other two comparisons (cRH vs. cLH and cRH vs. NcRH), the effect sizes that can be excluded are slightly larger |*g*| > 0.26 (ca. 1.5% explained variance). The meta-analyses of the sub-sectional data were comparatively less powerful resulting in wide confidence limits (see e.g., Fig. 5) as a result of the smaller number of studies and reduced overall sample size. Of note, however, also for the isthmus region, which had shown the largest difference between cRH and NcRH in Witelson’s original studies, effects larger |*g*| > 0.21 appear unlikely.

Thus, taken together, the present meta-analyses of differences in total corpus callosum size can be taken to exclude the existences of population effects larger than 1% or 1.5% explained variance with reasonable confidence. These effects would be considered “small” following Cohen’s effect-size conventions (Cohen, 1992). Thus, the population effect sizes that cannot be excluded may be seen as being irrelevant and handedness-related differences in corpus callosum size could be considered negligible for cognition and behaviour. On the other hand, as pointed out by Smith (2005), comparatively low amount of explained variances may have a substantial behavioural significance if they accumulate over many events. This may be the case with regard to the corpus callosum, considering its central role in the integration and coordination of information processing between the cerebral hemispheres. Thus, while the present analyses support the exclusion of comparatively small population effect sizes, we are hesitant to attributing functional insignificance to the effect size as it cannot be excluded. Any such conclusion would demand a better understanding of the relationship between variations in size and callosal functioning than currently available. Nevertheless, population-effect sizes of medium to large sizes, as were suggested by Witelson’s orginal work, can be excluded by the present analyses. Thus, we were not able to confirm the original findings neither in terms of the size of the effect nor in terms of statistical significance.

A closer inspection of the forest plots (Fig. 4 and 7) shows that the effect sizes of Witelson’s findings fall outside the 95% confidence interval of the estimated effect size for the respective meta-analysis as well as the prediction interval. The reasons explaining this strong deviation from the effects reported by other studies might be found in several characteristics of Witelson’s studies (Witelson, 1985, 1989; Witelson & Goldsmith, 1991). That is, the data was collected from autopsy specimen (rather than from in-vivo MRI), the study sample was comparatively old, and it was consisting only of (non-CNS) cancer patients. While each of these variables might affect callosal measurements, it appears, however, unlikely that these factors introduce systematic differences between right- and non-right handers. For example, while aging certainly has an effect on corpus callosum morphology (Danielsen et al., 2020; Doraiswamy et al., 1991; Hasan, Ewing-Cobbs, Kramer, Fletcher, & Narayana, 2008; Salat, Ward, Kaye, & Janowsky, 1997; Skumlien, Sederevicius, Fjell, Walhovd, & Westerhausen, 2018), the magnitude of aging-related callosal atrophy reduction would have to be more pronounced in NcRH individuals to produce the effect reported by Witelson. Likewise, cancer treatment (e.g., chemotherapy) negatively affects brain white-matter integrity, including the corpus callosum (Deprez et al., 2012), but to the best of our knowledge there is no evidence to suggest that NcRH are more resilient to these adverse side effects. An alternative explanation might be found when looking at the mean corpus callosum area reported across the three original studies. While the study sample was step-wise expanded by Witelson, the total corpus callosum area reported in particular for the male NcRH subsample decreased from a (comparatively high) value of 800.6 mm^2^ (sd: 53.9; *n* = 5) in 1985, to 786.3 mm^2^ (sd: 111.6; *n* = 6) in 1989, and to 744.0 mm^2^ (sd: 109.0; *n* = 9) in 1991. At the same time, all other subsample means stayed relatively stable. Therefore, one might speculate that outliers in the initial NcRH sample might have driven Witelson’s findings. Thus, Witelson’s finding of a larger corpus callosum in NcRH/MH might have been the consequence of a sampling bias.

Another issue that deserves attention is that the present null findings contrast a previous meta-analysis by Driesen and Raz (1995), which found a small effect size of Cohen’s *d* = −0.13 (CI 95%: −0.23 to −0.02), suggesting a significantly larger corpus callosum area in non-right handers. This meta-analysis included data from seven studies (Clarke & Zaidel, 1994; Kertesz, Polk, Howell, & Black, 1987; Nasrallah et al., 1986; O’Kusky et al., 1988; Steinmetz et al., 1992; Witelson, 1989; Yoshii & Duara, 1989), of which the present analyses included five (excluded were: O’Kusky et al., 1988, data only presented for patients with epilepsy, and Yoshii & Duara, 1989, published in Japanese). A closer look at the Driesen and Raz (1995) meta-analysis, however, also reveals some differences in the chosen approach. For example, the authors did not account for differences in handedness classification, and ignored, for example, that Nasrallah (1986) compares dRH and dLH participants, while others (Kertesz et al., 1987; Steinmetz et al., 1992; Witelson, 1989) compare cRH with NcRH samples. While this lack of specificity is certainly the consequence of the sparsity of studies available in 1995, which prevented a more sophisticated distinction, some inconsistencies in the data extraction remain difficult to explain. For example, in Table 3 (p. 243) Driesen and Raz list an effect size of Cohen’s *d* = 0.10 for the Steinmetz et al. (1992) study which would indicate larger areas in right-handers. Looking at the data provided in the original study, it rather is the case that the right-handers had the smallest corpus callosum compared to both MH and cLH groups. Furthermore, Driesen and Raz (1995) report an effect size of *d* = −0.57 for a study by O’Kusky et al. (1988). O’Kusky et al. do not report relevant group differences for the corpus callosum between left- and right-handers. The study only provides a correlation of *r* = −0.13 of handedness LI and callosal area for the studied sample of epileptic patients which might be converted to *d* = −0.26. Consequently, it is not immediately clear where the effect size used in the Driesen and Raz meta-analysis originated.

One limitation of the present meta-analyses is that we were not able to evaluate a handedness-related callosal differences controlling for brain-size differences, as the original studies typically did not consider brain size in their analyses. Thus, it can only be speculated how accounting for brain size might affect the handedness comparison. On the one hand, corpus callosum size and brain size are correlated positively (Jancke et al., 1997) so that possible brain-size differences between handedness groups might potentially act as a confounding variable. On the other hand, neither total brain volume nor white or gray matter volume have been found to differ between handedness groups in previous studies (Good et al., 2001; Hervé, Crivello, Perchey, Mazoyer, & Tzourio-Mazoyer, 2006; Jancke et al., 1997; Luders et al., 2010), making a systematic effect less likely. Arguably, most informative for this question are studies reporting analyses of both raw and brain-size corrected data allowing for a direct comparison. Three of these studies did not find any obvious differences for the handedness effect on total corpus callosum size when the correction is applied (Jancke et al., 1997; Mitchell et al., 2003; Nasrallah et al., 1986). Hopper et al. (1994) claim in a table legend that the body of the corpus callosum was found to be significantly larger in cRH compared to NcRH only after correction for brain size. However, given the currently available data, it appears unlikely that correcting corpus callosum for brain size, would substantial change the conclusions compared to the analysis of the raw callosal measures.

A second variable that needs to be considered is the participants’ sex, as it may act as both moderator and confounding variable. Sex may be considered a moderator variable, as a series of early studies report an interaction of sex and handedness when predicting total callosal and, in particular, isthmus size (Burke & Yeo, 1994; Clarke & Zaidel, 1994; Denenberg et al., 1991; Habib et al., 1991; Witelson, 1989). That is, a significantly larger isthmus area in NcRH compared to cRH has been found in male but not female participants (Habib et al., 1991; Witelson, 1989). Burke and Yeo (1994) utilizing the raw LQ score, report a positive correlation of the hand preference and posterior callosal area measures in their male subsample (suggesting that more consistent handedness is associated with a larger structure), while a negative association was reported in the female subsample. However, the present meta-analyses do neither for total nor isthmus area support the notion of an interaction. Firstly, across both meta-analyses neither males nor females showed a significant handedness effect for the cRH-NcRH comparison. Secondly, the direct comparison of the effect sizes did not yield a significant difference between males and females. Interpreting these findings, one has to keep in mind that the test power of the sex-specific and moderator analyses is reduced compared to omnibus analyses as fewer studies and smaller samples were included. Moreover, the power of moderator variables analysis itself is typically low within meta-analyses. Thus, arguably, more datasets would be beneficial to improve the meta-analyses’ sensitivity when it comes to the overall effect estimate, but even more so when it comes to the moderator variables analysis. Nevertheless, the overall pattern of findings provided no evidence for a strong interaction of sex and handedness when explaining corpus callosum morphology.

Meta-analyses and large scale studies usually find the corpus callosum to be comparable between the sexes or slightly smaller in males once differences in brain size have been accounted for (Driesen & Raz, 1995; Eliot, Ahmed, Khan, & Patel, 2021; Smith, 2005). For the here relevant uncorrected raw measures, however, the male corpus callosum can be expected to be bigger (Bishop & Wahlsten, 1997; Luders, Toga, & Thompson, 2014; Smith, 2005). Thus, differences in the proportions of male and female participants in the compared handedness groups might potentially bias the comparison and act as confounding variable. However, a look at the available data (see Tables S1 to S4), indicates that the proportions typically were matched well between the compared handedness groups. The study with the strongest differences in favour of NcRH by (Habib et al., 1991) even had more male participants in the NcRH than in cRH group, which should reverse the effect rather than enhance it. To further explore the possibility of a confounding influence of sex, we conducted supplementary meta-regression analyses using the difference in the proportion of females between handedness groups as a covariate. As can be seen in the Supplementary Material Section 9, for neither the dRH-dLH nor the cRH-NcRH comparison did we find a significant moderation effect. Thus, while differences in the proportions of males and females might affect the heterogeneity in the data, the differences in proportions are small and do not produce an obvious moderation effect, so that we do not believe that these have significantly affected the estimation of the mean effects.

Witelson emphasizes in her studies the relevance of consistency rather than the direction of hand preference (Witelson & Goldsmith, 1991). Best comparable with this approach is the meta-analysis comparing cRH and MH participants, which did not yield a significant effect. However, it also deserves to be mentioned that a small series of studies chose a slightly different approach to the topic by comparing a group of consistent hand preference – containing both cRH and cLH participants – with a non-consistent group (McDowell, Felton, Vazquez, & Chiarello, 2016; Welcome et al., 2009) or which used absolute LQ values in a correlative approach (Habib et al., 1991; Luders et al., 2010). While the group comparisons did not yield any significant association, Habib et al. (1991) found a significant positive correlation, indicating that more consistent individuals had a larger total corpus callosum area (Spearmann *r*_sp_ = 0.297, *p* = 0.03; *n* = 53). Luders et al. (2010), correlating callosal thickness measures with |LQ|, found a cluster of negative associations in the middle third of the corpus callosum (*n* =361, of which 37 were dLH), thus showing the opposite association to Habib et al. To further explore this association, we conducted a supplementary analysis of the samples of which raw data was available (Labache et al., 2020; McDowell et al., 2016; Van der Haegen, Cai, Seurinck, & Brysbaert, 2011; Westerhausen et al., 2004). As can been seen in Supplementary Material Section 10, the Spearman correlations of total corpus callosum area and |LQ| for these four samples ranged form *r*_sp_ = −0.15 to 0.10, with none of the correlations being significant. Although we refrained from conducting a meta-analysis for these studies, as their number was small, the available evidence does not support the existence of a substantial association of hand preference consistency and callosal measures.

Another consideration is that the present analysis focuses on morphological measures of the corpus callosum and will not be sensitive to differences on the microstructural level. While combined morphological and histological analyses suggest that the midsagittal area is a good predictor of the number of myelinated axons in the corpus callosum (Hou & Pakkenberg, 2012; Riise & Pakkenberg, 2011), studies comparing the diffusion characteristics of the corpus callosum between handedness groups could provide additional information. Three previous studies indeed suggest stronger anisotropy in the corpus callosum of non-right handers compared to right-handers (McKay, Iwabuchi, Häberling, Corballis, & Kirk, 2017; Westerhausen et al., 2004; Westerhausen et al., 2003); all comparing cRH and cLH), potentially suggesting a higher average axon or myelin density in the corpus callosum. Other studies, however, failed to find comparable differences (Haberling et al., 2011; Peled, Gudbjartsson, Westin, Kikinis, & Jolesz, 1998) revealing inconsistency in the literature. Unfortunately, the number of available studies is limited, so that we were not able to conduct a meta-analysis of diffusion data.

Handedness may be assessed via self-report preference questionnaires or via differences in left- and right-hand performance (skill) in manual tasks (Tapley & Bryden, 1985). As both not necessarily correlate highly with each other (Corey, Hurley, & Foundas, 2001), we originally had intention to perform a separate meta-analysis of studies utilizing measures of hand skill for the assessment of handedness. However, we were only able to identify three such studies so that we refrained from conducting a statistical integration. Kertesz et al. (1987) correlated performance differences between the right and the left hand in a tapping test with total corpus callosum size and did not find a significant association (*r* = 0.07, *N* = 104). Steinmetz, Staiger, Schlaug, Huang, and Jäncke (1995) used a paper-and-pencil manual tracing test to classify individuals into dLH (*n* = 58) and dRH individuals (*n* = 62), and did not find a significant difference in corpus-callosum size corrected for forebrain volume. Finally, Preuss et al. (2002), using the same manual tracing test as Steinmetz et al (1995), classified a group of nominal right-handed individuals into cRH (*n* = 32) and NcRH sample (*n* =14), and did not find a significant difference in total callosal size or in any subsection. Thus, while the available evidence does not indicate that corpus-callosum differences can be found when handedness is determined via hand-performance measures, the number of studies is small and more evidence is required before a conclusion can be reached.

Finally, the following potential biases from selective reporting and excluded studies deserve consideration when interpreting the findings. Firstly, for most of our meta-analyses concerning the total corpus callosum report bias is not indicated (through funnel plots and Egger’s regression analysis). Nevertheless, some studies, especially when analyzing subsectional data, selectively reported findings based on theoretical consideration (i.e., exclude regions from the analysis as previous studies did not find differences in these regions; e.g., Steinmetz et al., 1992; Witelson & Goldsmith, 1991) or statistical significance (Hines et al., 1992; Martens, Wilson, Chen, Wood, & Reutens, 2013; Moffat et al., 1998). In the latter case, this results in under reporting of non-significant, presumably small effect sizes, biasing the meta-analyses to overestimate the population-effect sizes (even when not noticeable in funnel plots). Thus, the effect-size estimates as shown in Fig 5 need to be interpreted with caution considering this bias. Secondly, data from several relevant publications had to be excluded as necessary information or data was missing (for details see Supplement Table S5) or not available even after contacting the authors. While the findings of several of these studies were included in the discussion, it has to be emphasized that most of these excluded studies indicate non-significant differences between handedness groups. Thus, it appears likely that the results of the present meta-analyses overestimate the size of the population-effect sizes. However, as none of the meta-analyses yielded significance in the first place, this overestimation of the effect size is unlikely to have affected the conclusions drawn from the present meta-analyses.

In summary, the general interpretation of Witelson’s findings (Witelson, 1985, 1989; Witelson & Goldsmith, 1991) that non-consistent handedness is associated with stronger inter-hemispheric connectivity, is not supported by the present meta-analyses of corpus callosum morphology. About 35 years after the first publication, the original findings have been rendered unreliable by the research they inspired. Thus, any assumption about callosal connections that is made based on hand preference are likely invalid. However, future large-scale studies are required to allow for a more powerful evaluation of sex differences, brains-size effects, as well as performance measures when studying handedness effects in the corpus callosum.

## Acknowledgments

We would like to thank the authors of the original publications who kindly provided the data and information required for the present meta-analyses. Only thanks to their effort, we were able to conduct meta-analysis of good statistical power. The HCP sample data was provided by the Human Connectome Project, WU-Minn Consortium (Principal Investigators: David Van Essen and Kamil Ugurbil; 1U54MH091657) funded by the 16 NIH Institutes and Centers that support the NIH Blueprint for Neuroscience Research; and by the McDonnell Center for Systems Neuroscience at Washington University.The project was supported by funding from the Department of Psychology, University of Oslo, to RW.

The references marked with an asterisk (*) were included the meta-analyses.

## References

Annett, M. (1970). A classification of hand preference by association analysis. British journal of psychology, 61(3), 303–321.

Beaton, A. A. (1997). The relation of planum temporale asymmetry and morphology of the corpus callosum to handedness, gender, and dyslexia: A review of the evidence. Brain and language, 60(2), 255–322.

Bishop, K. M., & Wahlsten, D. (1997). Sex differences in the human corpus callosum: myth or reality? Neurosci Biobehav Rev, 21(5), 581–601.

Bryden, M. (1977). Laterality functional asymmetry in the intact brain. New York, USA: Academic Press.

Budisavljevic, S., Castiello, U., & Begliomini, C. (2020). Handedness and White Matter Networks. The Neuroscientist, 1073858420937657.

Burke, H. L., & Yeo, R. A. (1994). Systematic variations in callosal morphology: The effects of age, gender, hand preference, and anatomic asymmetry. Neuropsychology, 8(4), 563–571.

Clarke, J. M., Lufkin, R. B., & Zaidel, F. (1993). Corpus-callosum morphometry and dichotic-listening perfromance - individual diiderences in functional interhemispheric inhibition. Neuropsychologia, 31(6), 547–557. doi:10.1016/0028-3932(93)90051-z

* Clarke, J. M., & Zaidel, E. (1994). Anatomical-behavioral relationships: corpus callosum morphometry and hemispheric specialization. Behav Brain Res, 64(1-2), 185–202. doi:10.1016/0166-4328(94)90131-7

Cohen, J. (1992). A power primer. Psychological bulletin, 112(1), 155.

Corey, D. M., Hurley, M. M., & Foundas, A. L. (2001). Right and left handedness defined: a multivariate approach using hand preference and hand performance measures. Cognitive and Behavioral Neurology, 14(3), 144–152.

* Cowell, P. E., & Gurd, J. (2018). Handedness and the Corpus Callosum: A Review and Further Analyses of Discordant Twins. Neuroscience, 388, 57–68. doi:10.1016/j.neuroscience.2018.06.017

Cowell, P. E., Kertesz, A., & Denenberg, V. H. (1993). Multiple dimensions of handedness and the human corpus callosum. Neurology, 43(11), 2353–2357. doi:10.1212/wnl.43.11.2353

Danielsen, V. M., Vidal-Piñeiro, D., Mowinckel, A. M., Sederevicius, D., Fjell, A. M., Walhovd, K. B., & Westerhausen, R. (2020). Lifespan trajectories of relative corpus callosum thickness: regional differences and cognitive relevance. Cortex, 130, 127–141.

* Denenberg, V. H., Kertesz, A., & Cowell, P. E. (1991). A factor analysis of the human’s corpus callosum. Brain Res, 548(1-2), 126–132. doi:10.1016/0006-8993(91)91113-f

Deprez, S., Amant, F., Smeets, A., Peeters, R., Leemans, A., Van Hecke, W., … Vandenbulcke, M. (2012). Longitudinal assessment of chemotherapy-induced structural changes in cerebral white matter and its correlation with impaired cognitive functioning. Journal of Clinical Oncology, 30(3), 274–281.

Doraiswamy, P. M., Figiel, G. S., Husain, M. M., McDonald, W. M., Shah, S. A., Boyko, O. B., … Krishnan, K. R. (1991). Aging of the human corpus callosum: magnetic resonance imaging in normal volunteers. J Neuropsychiatry Clin Neurosci, 3(4), 392–397. doi:10.1176/jnp.3.4.392

Driesen, N. R., & Raz, N. (1995). The influence of sex, age, and handedness on corpus callosum morphology: A meta-analysis. Psychobiology, 23(3), 240–247.

Eliot, L., Ahmed, A., Khan, H., & Patel, J. (2021). Dump the “dimorphism”: Comprehensive synthesis of human brain studies reveals few male-female differences beyond size. Neuroscience & Biobehavioral Reviews, 125, 667–697.

Friedrich, P., Fraenz, C., Schlüter, C., Ocklenburg, S., Mädler, B., Güntürkün, O., & Genç, E. (2020). The relationship between axon density, myelination, and fractional anisotropy in the human Corpus callosum. Cerebral Cortex, 30(4), 2042–2056.

Galaburda, A. M., Rosen, G. D., & Sherman, G. F. (1990). Individual variability in cortical organization: its relationship to brain laterality and implications to function. Neuropsychologia, 28(6), 529–546.

Gazzaniga, M. S. (2000). Cerebral specialization and interhemispheric communication: does the corpus callosum enable the human condition? Brain, 123 (Pt 7), 1293–1326.

Good, C. D., Johnsrude, I., Ashburner, J., Henson, R. N., Friston, K. J., & Frackowiak, R. S. (2001). Cerebral asymmetry and the effects of sex and handedness on brain structure: a voxel-based morphometric analysis of 465 normal adult human brains. Neuroimage, 14(3), 685–700.

Gurd, J. M., Cowell, P. E., Lux, S., Rezai, R., Cherkas, L., & Ebers, G. C. (2013). fMRI and corpus callosum relationships in monozygotic twins discordant for handedness. Brain Structure & Function, 218(2), 491–509. doi:10.1007/s00429-012-0410-9

* Haberling, I. S., Badzakova-Trajkov, G., & Corballis, M. C. (2011). Callosal tracts and patterns of hemispheric dominance: A combined fMRI and DTI study. Neuroimage, 54(2), 779–786. doi:10.1016/j.neuroimage.2010.09.072

* Haberling, I. S., Badzakova-Trajkov, G., & Corballis, M. C. (2012). The Corpus Callosum in Monozygotic Twins Concordant and Discordant for Handedness and Language Dominance. Journal of Cognitive Neuroscience, 24(10), 1971–1982. Retrieved from <Go to ISI>://WOS:000308422200001

* Habib, M., Gayraud, D., Oliva, A., Regis, J., Salamon, G., & Khalil, R. (1991). Effects of handedness and sex on the morphology of the corpus callosum: a study with brain magnetic resonance imaging. Brain Cogn, 16(1), 41–61.

Harrer, M., Cuijpers, P., Furukawa, T., & Ebert, D. D. (2019). dmetar: Companion R Package For The Guide ‘Doing Meta-Analysis in R’. Retrieved from http://dmetar.protectlab.org/

Hasan, K. M., Ewing-Cobbs, L., Kramer, L. A., Fletcher, J. M., & Narayana, P. A. (2008). Diffusion tensor quantification of the macrostructure and microstructure of human midsagittal corpus callosum across the lifespan. NMR Biomed, 21(10), 1094–1101. doi:10.1002/nbm.1286

Hervé, P.-Y., Crivello, F., Perchey, G., Mazoyer, B., & Tzourio-Mazoyer, N. (2006). Handedness and cerebral anatomical asymmetries in young adult males. Neuroimage, 29(4), 1066–1079.

* Hines, M., Chiu, L., McAdams, L. A., Bentler, P. M., & Lipcamon, J. (1992). Cognition and the corpus callosum: verbal fluency, visuospatial ability, and language lateralization related to midsagittal surface areas of callosal subregions. Behavioral neuroscience, 106(1), 3.

Hopper, K. D., Patel, S., Cann, T. S., Wilcox, T., & Schaeffer, J. M. (1994). The relationship of age, gender, handedness, and sidedness to the size of the corpus callosum. Acad Radiol, 1(3), 243–248. doi:10.1016/s1076-6332(05)80723-8

Hou, J., & Pakkenberg, B. (2012). Age-related degeneration of corpus callosum in the 90+ years measured with stereology. Neurobiol Aging, 33(5), 1009 e1001–1009. doi:10.1016/j.neurobiolaging.2011.10.017

* Jancke, L., Staiger, J. F., Schlaug, G., Huang, Y. X., & Steinmetz, H. (1997). The relationship between corpus callosum size and forebrain volume. Cerebral Cortex, 7(1), 48–56. doi: 10.1093/cercor/7.1.48

Josse, G., Seghier, M. L., Kherif, F., & Price, C. J. (2008). Explaining Function with Anatomy: Language Lateralization and Corpus Callosum Size. Journal of Neuroscience, 28(52), 14132–14139. doi:10.1523/jneurosci.4383-08.2008

Karolis, V. R., Corbetta, M., & De Schotten, M. T. (2019). The architecture of functional lateralisation and its relationship to callosal connectivity in the human brain. Nature communications, 10(1), 1–9.

* Kertesz, A., Polk, M., Howell, J., & Black, S. E. (1987). Cerebral dominance, sex, and callosal size in MRI. Neurology, 37(8), 1385–1388. doi:10.1212/wnl.37.8.1385

* Labache, L., Mazoyer, B., Joliot, M., Crivello, F., Hesling, I., & Tzourio-Mazoyer, N. (2020). Typical and atypical language brain organization based on intrinsic connectivity and multitask functional asymmetries. Elife, 9. doi:10.7554/eLife.58722

Lüdecke, D. (2018). esc: Effect Size Computation for Meta Analysis. Retrieved from https://CRAN.R-project.org/package=esc

Luders, E., Cherbuin, N., Thompson, P. M., Gutman, B., Anstey, K. J., Sachdev, P., & Toga, A. W. (2010). When more is less: associations between corpus callosum size and handedness lateralization. Neuroimage, 52(1), 43–49. doi:10.1016/j.neuroimage.2010.04.016

* Luders, E., Rex, D. E., Narr, K. L., Woods, R. P., Jancke, L., Thompson, P. M., … Toga, A. W. (2003). Relationships between sulcal asymmetries and corpus callosum size: Gender and handedness effects. Cerebral Cortex, 13(10), 1084–1093. doi:10.1093/cercor/13.10.1084

Luders, E., Toga, A. W., & Thompson, P. M. (2014). Why size matters: differences in brain volume account for apparent sex differences in callosal anatomy: the sexual dimorphism of the corpus callosum. Neuroimage, 84, 820–824. doi:10.1016/j.neuroimage.2013.09.040

* Martens, M. A., Wilson, S. J., Chen, J., Wood, A. G., & Reutens, D. C. (2013). Handedness and corpus callosal morphology in Williams syndrome. Development and Psychopathology, 25(1), 253–260. doi:10.1017/s0954579412001009

* McDowell, A., Felton, A., Vazquez, D., & Chiarello, C. (2016). Neurostructural correlates of consistent and weak handedness. Laterality, 21(4-6), 348–370. doi:10.1080/1357650x.2015.1096939

McKay, N. S., Iwabuchi, S. J., Häberling, I. S., Corballis, M. C., & Kirk, I. J. (2017). Atypical white matter microstructure in left-handed individuals. Laterality: Asymmetries of Body, Brain and Cognition, 22(3), 257–267.

Mitchell, T. N., Free, S. L., Merschhemke, M., Lemieux, L., Sisodiya, S. M., & Shorvon, S. D. (2003). Reliable callosal measurement: Population normative data confirm sex-related differences. American Journal of Neuroradiology, 24(3), 410–418.

* Moffat, S. D., Hampson, E., & Lee, D. H. (1998). Morphology of the planum temporale and corpus callosum in left handers with evidence of left and right hemisphere speech representation. Brain, 121, 2369–2379. doi:10.1093/brain/121.12.2369

* Morton, B. E., & Rafto, S. E. (2006). Corpus callosum size is linked to dichotic deafness and hemisphericity, not sex or handedness. Brain and cognition, 62(1), 1–8. doi:10.1016/j.bandc.2006.03.001

* Nasrallah, H. A., Andreasen, N. C., Coffman, J. A., Olson, S. C., Dunn, V. D., Ehrhardt, J. C., & Chapman, S. M. (1986). A controlled magnetic resonance imaging study of corpus callosum thickness in schizophrenia. Biol Psychiatry, 21(3), 274–282. doi:10.1016/0006-3223(86)90048-x

O’Kusky, J., Strauss, E., Kosaka, B., Wada, J., Li, D., Druhan, M., & Petrie, J. (1988). The corpus callosum is larger with right-hemisphere cerebral speech dominance. Annals of Neurology: Official Journal of the American Neurological Association and the Child Neurology Society, 24(3), 379–383.

Ocklenburg, S., & Güntürkün, O. (2018). The lateralized brain: The neuroscience and evolution of hemispheric asymmetries. London, UK: Academic Press.

Oldfield, R. C. (1971). The assessment and analysis of handedness: the Edinburgh inventory. Neuropsychologia, 9(1), 97–113.

* Ozdikici, M. (2020). Measurment of midsagittal corpus callosum area with the modified Cavalieri method in healthy right- and left-handed Turkish adults. Malang Neurology Journal, 6(1), 24–27.

Page, M. J., Moher, D., Bossuyt, P. M., Boutron, I., Hoffmann, T. C., Mulrow, C. D., … Brennan, S. E. (2021). PRISMA 2020 explanation and elaboration: updated guidance and exemplars for reporting systematic reviews. bmj, 372, n160.

Papadatou-Pastou, M., Ntolka, E., Schmitz, J., Martin, M., Munafò, M. R., Ocklenburg, S., & Paracchini, S. (2020). Human handedness: A meta-analysis. Psychological bulletin, 146(6), 481–524.

Parker, A., Parkin, A., & Dagnall, N. (2017). Effects of handedness & saccadic bilateral eye movements on the specificity of past autobiographical memory & episodic future thinking. Brain and cognition, 114, 40–51.

Peled, S., Gudbjartsson, H., Westin, C.-F., Kikinis, R., & Jolesz, F. A. (1998). Magnetic resonance imaging shows orientation and asymmetry of white matter fiber tracts. Brain research, 780(1), 27–33.

Preuss, U. W., Meisenzahl, E. M., Frodl, T., Zetzsche, T., Holder, J., Leinsinger, G., … Möller, H. J. (2002). Handedness and corpus callosum morphology. Psychiatry Res, 116(1-2), 33–42. doi:10.1016/s0925-4927(02)00064-1

Prichard, E., Propper, R. E., & Christman, S. D. (2013). Degree of handedness, but not direction, is a systematic predictor of cognitive performance. Frontiers in psychology, 4, 9.

Riise, J., & Pakkenberg, B. (2011). Stereological estimation of the total number of myelinated callosal fibers in human subjects. Journal of Anatomy, 218(3), 277–284.

Ringo, J. L., Doty, R. W., Demeter, S., & Simard, P. Y. (1994). Time is of the essence: a conjecture that hemispheric specialization arises from interhemispheric conduction delay. Cerebral Cortex, 4(4), 331–343.

Roberts, B. R., Fernandes, M. A., & MacLeod, C. M. (2020). Re-evaluating whether bilateral eye movements influence memory retrieval. PLoS One, 15(1), e0227790.

Sala, G., Signorelli, M., Barsuola, G., Bolognese, M., & Gobet, F. (2017). The relationship between handedness and mathematics is non-linear and is moderated by gender, age, and type of task. Frontiers in psychology, 8, 948.

Salat, D., Ward, A., Kaye, J. A., & Janowsky, J. S. (1997). Sex differences in the corpus callosum with aging. Neurobiol Aging, 18(2), 191–197. doi:10.1016/s0197-4580(97)00014-6

Schwarzer, G. (2020). meta: General Package for Meta-Analysis. Retrieved from https://CRAN.R-project.org/package=meta

Skumlien, M., Sederevicius, D., Fjell, A. M., Walhovd, K. B., & Westerhausen, R. (2018). Parallel but independent reduction of emotional awareness and corpus callosum connectivity in older age. PLoS One, 13(12), e0209915.

Smith, R. J. (2005). Relative size versus controlling for size. Current Anthropology, 46(2), 249–273.

* Steinmetz, H., Jäncke, L., Kleinschmidt, A., Schlaug, G., Volkmann, J., & Huang, Y. (1992). Sex but no hand difference in the isthmus of the corpus callosum. Neurology, 42(4), 749–752. doi:10.1212/wnl.42.4.749

Steinmetz, H., Staiger, J. F., Schlaug, G., Huang, Y., & Jäncke, L. (1995). Corpus callosum and brain volume in women and men. Neuroreport, 6(7), 1002–1004. doi:10.1097/00001756-199505090-00013

Tapley, S., & Bryden, M. (1985). A group test for the assessment of performance between the hands. Neuropsychologia, 23(2), 215–221.

* Tuncer, M. C., Hatipoglu, E. S., & Ozates, M. (2005). Sexual dimorphism and handedness in the human corpus callosum based on magnetic resonance imaging. Surg Radiol Anat, 27(3), 254–259. doi:10.1007/s00276-004-0308-1

* Van der Haegen, L., Cai, Q., Seurinck, R., & Brysbaert, M. (2011). Further fMRI validation of the visual half field technique as an indicator of language laterality: A large-group analysis. Neuropsychologia, 49(10), 2879–2888.

* Van Essen, D. C., Smith, S. M., Barch, D. M., Behrens, T. E., Yacoub, E., Ugurbil, K., & HCP-Consortium. (2013). The WU-Minn human connectome project: an overview. Neuroimage, 80, 62–79.

Viechtbauer, W. (2020). metafor: Meta-Analysis Package for R. Retrieved from https://CRAN.R-project.org/package=metafor

Welcome, S. E., Chiarello, C., Towler, S., Halderman, L. K., Otto, R., & Leonard, C. M. (2009). Behavioral correlates of corpus callosum size: Anatomical/behavioral relationships vary across sex/handedness groups. Neuropsychologia, 47(12), 2427–2435. doi: 10.1016/j.neuropsychologia.2009.04.008

* Westerhausen, R., Kreuder, F., Dos Santos Sequeira, S. D., Walter, C., Woerner, W., Wittling, R. A., … Wittling, W. (2004). Effects of handedness and gender on macro- and microstructure of the corpus callosum and its subregions: a combined high-resolution and diffusion-tensor MRI study. Cognitive Brain Research, 21(3), 418–426. doi:10.1016/j.cogbrainres.2004.07.002

Westerhausen, R., Kreuder, F., Sequeira, S. D. S., Walter, C., Woerner, W., Wittling, R. A., … Wittling, W. (2006). The association of macro-and microstructure of the corpus callosum and language lateralisation. Brain and language, 97(1), 80–90.

Westerhausen, R., Walter, C., Kreuder, F., Wittling, R. A., Schweiger, E., & Wittling, W. (2003). The influence of handedness and gender on the microstructure of the human corpus callosum: a diffusion-tensor magnetic resonance imaging study. Neuroscience letters, 351(2), 99–102.

* Witelson, S. F. (1985). The brain connection: the corpus callosum is larger in left-handers. Science, 229(4714), 665–668. doi:10.1126/science.4023705

* Witelson, S. F. (1989). Hand and sex differences in the isthmus and genu of the human corpus callosum. A postmortem morphological study. Brain, 112 (Pt 3), 799–835. doi:10.1093/brain/112.3.799

* Witelson, S. F., & Goldsmith, C. H. (1991). The relationship of hand preference to anatomy of the corpus callosum in men. Brain Res, 545(1-2), 175–182. doi:10.1016/0006-8993(91)91284-8

Witelson, S. F., & Nowakowski, R. S. (1991). Left out axons make men right: A hypothesis for the origin of handedness and functional asymmetry. Neuropsychologia, 29(4), 327–333.

Yoshii, F., & Duara, R. (1989). Size of the corpus callosum in normal subjects and patients with Alzheimer’s disease--magnetic resonance imaging study. Rinsho shinkeigaku= Clinical neurology, 29(1), 1–7.

Zapała, D., Zabielska-Mendyk, E., Augustynowicz, P., Cudo, A., Jaśkiewicz, M., Szewczyk, M., … Francuz, P. (2020). The effects of handedness on sensorimotor rhythm desynchronization and motor-imagery BCI control. Scientific reports, 10(1), 1–11.

