## Supplement Material for "Handedness and midsagittal corpus callosum morphology: A systematic meta-analytic evaluation"

Supplement information

### 1. Table S1 to S4. Studies included in meta-analyses with details regarding sample characteristics and data extraction.

*Table S1*. Studies included in the meta-analysis comparing dRH and dLH participants (a)

| Study | | Descriptive Statistics | | Comments | |
|---|---|---|---|---|---|
| # | Reference | %female(dRH/dLH) | Age | Data extraction | Other |
| **1** | Denenberg et al. (1991) | 52/50 | mean: 26.9 (7.2) yrs; range: 18 to 49 yrs | Data of total corpus callosum (tcc) and subsections average of male and female samples; see Table II (p. 128) and III (p.129) of original publication; subsection transformed following heuristics in Supplement Section 4 | Handedness info (a), and MRI field strength from Keretsz et al. (1987); total cc data is based on area measures, subsection data is based on thickness measures |
| **2** | Haberling et al. (2011) | 51/48 | mean: 20.8/23.2 (4.6/4.6) yrs (m/f); range: n.a. | Group means for tcc provided by author (see Supplement Section 3). | |
| **3** | Haberling et al. (2012) | 60 (b) | mean: 24.7 (8.8) yrs; range: 15 to 49 yrs | Group means for tcc provided by author (see Supplement Section 3). | The sample consists of monozygotic twins, concordant or discordant for hand preference. |
| **4** | HCP (Van Essen et al., 2013) | 55/47 | mean: 28.8 (3.7) yrs; range: 22 to 37 yrs | Raw data available and re-analysed (see Supplement Section 3) | |
| **5** | Labache et al. (2020) | 47/48 | mean: 25.9(6.5) yrs; range: 18 to 57 yrs | Raw data available and re-analysed (see Supplement Section 3) | |
| **6** | Martens et al. (2013) | n.a. | mean: 18.3 (11.4) yrs; range: 8 to 41 yrs | Control group data, Table 1 (p. 257) | Thickness data was not useable for integration; for subsections only midbody data presented; |
| **7** | McDowell et al. (2016) | 50/50(c) | mean: 21.4 (n.a.) yrs; range: n.a. | Raw data available and re-analysed (see Supplement Section 3); subsection transformed following heuristic in Supplement Section 4 | |

| | | | | | |
|---|---|---|---|---|---|
| **8** | Moffat et al. (1998) | 0/0 | mean: 24.5/23.3 (n.a.) yrs (cRH/cLH) | Data extraction: t values for tcc and two subsections comparison used (p. 2373); CC4 data used as mid third; CC5 as isthmus | |
| **9** | Morton and Rafto (2006) | 48 (b) | mean: 45.3 (14.2) yrs; range: 18 to 74 yrs | Data from Table 1, p.4 | |
| **10** | Nasrallah et al. (1986) | 0/0 | mean 28.0 (4.9) yrs; range: n.a. | Control group data, as reported in Tables 2 (p. 277) and 3 (p. 278) | Only data of male sample presented in article |
| **11** | Ozdikici (2020) | 57/57 | mean: n.a.; range: 20 to 69 yrs | Data average of male and female sample (see Table 2, p.3) | Author confirmed that writing and eating hand was used to determine hand preference |
| **12** | Tuncer et al. (2005) | 50/50 | mean: n.a.; range: 21 to 32 yrs | As reported on p. 256 in text | subsection data was not reported although analysed |
| **13** | Van der Haegen et al. (2011) | 83/75 | mean: 21.7 (4.3) yrs; range: 17 to 38 yrs | Raw data available and re-analysed (see Supplement Section 3) | CC data not previously published |
| **14** | Westerhausen et al. (2004) | 54/55 | mean: 24.0 (3.5) yrs; range: 19 to 38 yrs | Raw data available and re-analysed (see Supplement Section 3) | sample larger than described in original publication |

Notes. n.a. = not available, a) classification criterion for all studies was a LQ larger or below zero for the dRH and the dLH group, respectively; or preference in self-identification (study #9); b) in total sample; c) as original sample slightly differs in the number of subjects from the data available

*Table S2*. Studies included in the meta-analysis comparing cRH and NcRH participants

| Study | | Descriptive statistics | | | Comments | |
|---|---|---|---|---|---|---|
| # | Reference | HD criterion (a) | %female | Age | Data extraction | Other |
| **1** | Clarke et al. (1994) | Qualitative (a) | 50/50 (b) | mean: 28.2 (n.a.) yrs; range: 21 to 43 yrs | Data from Table 1 (p. 190; original publication); pooled across female and male samples; subsection: mid third subsection as pooled data of anterior and posterior midbody | |
| **2** | Cowell & Gurd (2018) | 70% of max LQ | 100/100 | mean: n.a.; range: 37 to 67 yrs | Data from Table 1 (p.61); subsection transformed following heuristics in Supplement Section 4 | The sample consists of monozygotic twins discordant for hand preference; tcc data is based on area measures, subsection data on thickness measures |
| **3** | Haberling et al. (2011) | cRH: ≥+80% LQ; NcRH: <+80% LQ | n.a. | mean: 20.8/23.2 (4.6/4.6) yrs (m/f); range: n.a. | Group means for tcc provided by author (see Supplement Section 3) | |
| **4** | Haberling et al. (2012) | cRH: ≥+80% LQ; NcRH: <+80% LQ | 60 (c) | mean: 24.7 (8.8) yrs; range: 15 to 49 yrs | Group means for tcc provided by author (see Supplement Section 3) | The sample consists of monozygotic twins, concordant or discordant for hand preference |
| **5** | Habib et al. (1991) | cRH: ≥+80% LQ; NcRH: <+80% LQ | 27/41 | mean: 29 (n.a.) yrs; range: 18 to 51 yrs | Data from Table 1 (p. 48); subsection transformed following heuristic in Supplement Section 4 | |
| **6** | HCP (Van Essen et al., 2013) | cRH: ≥+80% LQ; NcRH: <+80% LQ | 63/44 | mean: 28.8 (3.7.) yrs; range: 22 to 37 yrs | Raw data available and re-analysed (see Supplement Section 3) | |
| **7** | Hines et al. (1992) | Qualitative | 100/100 | mean: 29.4 (5.6) yrs range: 20 to 45 yrs | Only group means for isthmus provided (p. 6) | |
| **8** | Jäncke et al. (1997) | Qualitative | 35/45 | mean: 25.7 (4.7) yrs; range: 18 to 45 yrs | Data from Table 4 (p.51); NcRH data pooled average of "CLH" and "MH" groups | |
| **9** | Labache et al. (2020) | cRH: ≥+80% LQ; NcRH: <+80% LQ | 49/49 | mean: 25.8(6.5) yrs; range: 18 to 57 yrs | Raw data available and re-analysed (see Supplement Section 3) | |

| | | | | | | |
|---|---|---|---|---|---|---|
| **10** | McDowell et al. (2016) | cRH: ≥+80% LQ;<br>NcRH: <+80% LQ | 54/40 | n.a. | Raw data available and re-analysed (see Supplement Section 3); subsection transformed following heuristic in Supplement Section 4) | As the sample reported in the publication slightly differs from the data provided, it is not possible to determine the exact sex distribution or age |
| **11** | Steinmetz et al. (1992) | Qualitative | 47/52 | mean: 26.3 (n.a.) yrs;<br>range: 21 to 35 yrs | Data from Table on p. 751; NcRH data pooled average of "CLH" and "MH" group | Only data of posterior subsections reported in article |
| **12** | Van der Haegen (2011) | cRH: ≥+80% LQ;<br>NcRH: <+80% LQ | 79/77 | mean: 21.7 (4.3) yrs;<br>range: 17 to 38 yrs | Raw data available and re-analysed (see Supplement Section 3) | |
| **13** | Westerhausen et al. (2004) | cRH: ≥+80% LQ;<br>NcRH: <+80% LQ | 49/52 | mean: 24.0 (3.5) yrs;<br>range: 19 to 38 yrs | Raw data available and re-analysed (see Supplement Section 3) | |
| **14** | Witelson (1989); Witelson and Goldsmith (1991) | Qualitative | 64/57 | mean: 50.8 yrs;<br>range: 25 to 68 yrs | Pooled dat : female data from Witelson (1989, Table 3, p. 809); male tcc and isthmus data from Witelson & Goldsmith (1991, Table II, p. 177); subsections averaged where necessary | Sample consists of cancer patients |

*Notes.* n.a. = not available, a) handedness group classification criterion: "qualitative" indicates that it was based on the answer per individual item not on the LQ score respectively; b) where available %female is given by group (cRH/NcRH); c) for whole sample

*Table S3*. Studies included in the meta-analysis comparing cRH and cLH participants

| Study | | Descriptive statistics | | | Comments | |
|---|---|---|---|---|---|---|
| # | First author | Criterion (a) | %female | Age | Data extraction | Other |
| **1** | Haberling et al. (2011) | cRH: ≥+80% LQ; cLH: ≤-80% LQ | n.a. | n.a. | Group means for tcc provided by author (see Supplement Section 3) | |
| **2** | Haberling et al. (2012) | cRH: ≥+80% LQ; cLH: ≤-80% LQ | n.a. | n.a. | Group means for tcc provided by author (see Supplement Section 3) | The sample consists of monozygotic twins, concordant or discordant for hand preference. |
| **3** | HCP (Van Essen et al., 2013) | cRH: ≥+80% LQ; cLH: ≤-80% LQ | 63/64 | mean: 29.0 (3.6) yrs; range: 22 to 36 yrs | Raw data available and re-analysed (see Supplement Section 3) | |
| **4** | Jäncke et al. (1997) | Qualitative (b) | 35/54 (b) | n.a. | Data from Table 4 (p. 51, original publications) | |
| **5** | Kertesz et al. (1987) | cRH: ≥+50% LQ; cLH: ≤-50% LQ | n.a. | n.a. | Data from Table 1 (p. 1387); | |
| **6** | Labache et al. (2020) | cRH: ≥+80% LQ; cLH: ≤-80% LQ | 49/47 | mean: 26.3 (6.3) yrs; range: 18 to 53 yrs | Raw data available and re-analysed (see Supplement Section 3) | |
| **7** | Luders et al. (2003) | "strong LQs" (but cut-off not defined explicitely) | 50/47 | n.a. | Data from Table 1 (p. 1087); mid third calculated as mean of anterior and posterior midbody | |
| **8** | McDowell et al. (2016) | cRH: ≥+80% LQ; cLH: ≤-80% LQ | 54/56 | mean: 21.4 (n.a.) yrs; range: n.a. | Raw data available and reanalysed (see Supplement Section 3); subsection transformed following heuristic in Supplement Section 4 | |
| **9** | Steinmetz et al. (1992) | Qualitative | 47/44 | n.a. | Data from Table on p. 751 | Only data of posterior subsections reported in article |
| **10** | Van der Haegen (2011) | cRH: ≥+80% LQ; cLH: ≤-80% LQ | 79/79 | mean: 22.2 (4.4) yrs; range: 17 to 38 yrs | Raw data available and re-analysed (see Supplement Section 3) | |
| **11** | Westerhausen et al (2004) | cRH: ≥+80% LQ; cLH: ≤-80% LQ | 49/56 | mean: 24.2 (3.8) yrs; range: 19 to 38 yrs | Raw data available and re-analysed (see Supplement Section 3) | |

*Notes.* n.a. = not available, a) handedness group classification criterion: “qualitative” indicates that it was based on the answer per individual item not on the LQ score respectively; b) where available %female is given by group (cRH/cLH);

*Table S4*. Studies included in the meta-analysis comparing cRH and MH participants

| Study | | Descriptive statistics | | | Comments | |
|---|---|---|---|---|---|---|
| # | First author | Criterion (a) | %female | Age | Data extraction | Comments |
| **1** | Haberling et al. (2011) | cRH: ≥+80% LQ;<br>MH: btw. +80% and -80% LQ | n.a. | n.a. | Group means for tcc provided by author (see Supplement Section 3) | |
| **2** | Haberling et al. (2012) | cRH: ≥+80% LQ;<br>MH: btw. +80% and -80% LQ | n.a. | n.a. | Group means for tcc provided by author (see Supplement Section 3) | The sample consists of monozygotic twins, concordant or discordant for hand preference. |
| **3** | HCP (Van Essen et al., 2013) | cRH: ≥+80% LQ;<br>MH: btw. +80% and -80% LQ | 63/43 | mean: 28.8 (3.7) yrs;<br>range: 22 to 37 yrs | Raw data available and re-analysed (see Supplement Section 3) | |
| **4** | Jäncke et al. (1997) | Qualitative | 35/39(b) | n.a. | Data from Table 4 (p. 51, of original publication) | |
| **5** | Kertesz et al. (1987) | cRH: ≥+50% LQ;<br>MH: btw. +40% and -50/-60% LQ | n.a. | n.a. | Data from Table 1 (p. 1387) | Inconsistency in defining MH lower boundary (either -50% or -60% LQ), see p. 1386/1387 |
| **6** | Labache et al. (2020) | cRH: ≥+80% LQ;<br>MH: btw +80% and -80% LQ | 49/50 | mean: 26.3 (6.6) yrs;<br>range: 18 to 57 yrs | Raw data available and re-analysed (see Supplement Section 3) | |
| **7** | McDowell et al. (2016) | cRH: ≥+80% LQ;<br>MH: btw +80% and -80% LQ | 55/36 | mean: 21.4 (n.a.) yrs;<br>range: n.a. | Raw data available and reanalysed (see Supplement Section 3); subsection transformed following heuristic in Supplement Section 4 | |
| **8** | Steinmetz et al. (1992) | Qualitative | 47/54 | n.a. | Data from Table on p. 751 | Only data of posterior subsections reported in article |
| **9** | Van der Haegen (2011) | cRH: ≥+80% LQ;<br>MH: btw +80% and -80% LQ | 79/75 | mean: 22.2 (4.5) yrs;<br>range: 17 to 38 yrs | raw data available and reanalysed (see Supplement Section 3) | |
| **10** | Westerhausen et al (2004) | cRH: ≥+80% LQ; MH: btw +80% and -80% LQ | 49/43 | mean: 23.7 (2.7) yrs;<br>range: 19 to 33 yrs | raw data available and reanalysed (see Supplement Section 3) | |
| **11** | Witelson (1985) | Qualitative | 74/67 | mean: n.a. (n.a.) yrs;<br>range: 25 to 60 yrs | Data from Table 1 (p. 666) | |

*Notes.* n.a. = not available, a) handedness group classification criterion: “qualitative” indicates that it was based on the answer per individual item not on the LQ score respectively; b) where available %female is given by group (cRH/MH);

### 2. Table S5. Studies excluded from meta-analysis and reason for exclusion

*Table S5* Studies not included in the meta-analysis but which were considered eligible

| # | Study | Approach and narrative findings | Reason for exclusion |
|---|---|---|---|
| 1 | Bruke & Yeo (1994) | In men, posterior corpus callosum was correlated positively with increasing right-handedness (LQ score). In women, greater right-handedness was associated with smaller callosal areas in anterior and posterior corpus callosum. | Does not report group means, so that the study could not be included in the statistical analysis. |
| 2 | Cherbuin et al. (2013) | Analysis the relationship of brain asymmetries and corpus callosum thickness including a group of non-right handers. Handedness effects on callosal measures are, however, not explicitly reported. | Sample overlap with Luders et al. (2010); not included in the meta-analysis (see below) |
| 3 | Clarke et al. (1993) | Focusses on the relationship of corpus callosum morphology and dichotic listening performance. Differences between left- and right-handers in total corpus callosum area are reported as non-significant. | More complete report of results in Clarke and Zaidel (1994), so that data reported in Clarke et al. (1993) was considered redundant |
| 4 | Cowell et al. (1993) | Provides a more detailed analysis of the relationship of consistency and direction of hand preference and the corpus callosum, as well as an analysis of "throwing hand" as important factor. | Same sample as reported in other included studies (Denenberg et al., 1991; Kertesz et al., 1987); the data of Cowell et al. (1993) was thus redundant to the previous publications |
| 5 | Gurd et al. (2013) | Comparison between right- and left-handed twins non-significant except for the region W22–39 (approx. genu) with a larger average thickness in the LH twins. | The results of the extended sample reported in Cowell and Gurd (2018) was included in the meta-analyses; Gurd et al. (2013) was not included. |
| 6 | Hopper et al. (1994) | Report larger area of the anterior body of the corpus callosum in cRH compared to NcRH after correction for brain size. Comparison without correction was not significant. | The authors do not provide any measure of dispersion or test statistics so that the data cannot be included in the meta-analysis. |
| 7 | Josse et al. (2008) | In the supplement to the publication the authors report a non-significant association of the EHI test score with total corpus callosum area in multiple regressions analysis including sex, age, and hemispheric white matter volume. The association was significant in a subsection covering genu and anterior body of the corpus callosum, but in no other subsection. | The data was not included as the mean raw data was not available for handedness groups. |
| 8 | Luders et al. (2010) | The authors report negative correlations between callosal thickness and absolute Annett handedness scores in anterior and posterior midbody. Group comparisons between cRH and other handedness groups not significant. All analyses uses brain size as covariate. | Not included as uncorrected mean data for groups could not be obtained. |

| | | | |
|---|---|---|---|
| 9 | Mitchell et al. (2003) | Reports normative corpus callosum data. No significant correlations were noted between handedness and callosal area or the area:cerebral volume ratio. | Not included in meta-analysis as mean data not reported. |
| 10 | O'Kusky et al (1988) | Analyses focusses on differences between lateralisation groups, not handedness. Association of corpus callosum and handedness measures only reported for patients with epilepsy not for the control sample. | Not included in meta-analysis as data of healthy sample to presented. . |
| 11 | Preuss et al. (2002) | No difference in total or subsection area of the corpus callosum between consistent and non-consistent right handers. | All participants are nominal right-handers (dRH); classification is based on hand-skill/performance data, and the study was consequently not included in the meta-analysis. |
| 12 | Reinarz et al. (1988) | No differences between handedness groups (dRH vs dLH) in callosal area relative to brain size or in subsectional area relative to total corpus callosum size. | Only the subsection area relative to total corpus callosum area available in the article. Thus, the data cannot be included in the meta-analysis. |
| 13 | Robichon & Habib (1998) | Focus on dyslexia but includes 2 NcRH control participants. | Data presented in graph and cannot be extracted. |
| 14 | Steinmetz et al. (1995) | No difference between handedness groups in total corpus callosum area before or after adjustment for brain volume. | Handedness was defined based on hand performance, and is thus not included in the meta-analysis. However, a re-analysis of the data based on hand preference is reported by Jäncke et al (1997), which is included in the meta-analysis. |
| 15 | Welcome et al. (2009) | Report no differences between consistent handers (combined cRH/cLH) and participants of inconsistent preference after correcting for brain size. | Data provided was not sufficient for inclusion in the meta analysis. However, the data was presented after re-analysis by McDowell et al. (2016), and raw data of this study was included in the present meta-analysis. |
| 16 | Westerhausen et al. (2006) | Analysis based on language lateralisaton with handedness as covariate. | Sample is an extension of the Westerhausen et al. (2004) report and raw data of this study was included in the present meta-analysis. |

### 3. Extraction of data available only through author contact

This section reports the data that became available after contacting the authors and explains how the mean and standard deviations included in the meta-analyses of the total corpus callosum size were determined. The means and standard deviations for the subregions can be found in the data tables provided on the accompanying OSF platform (https://osf.io/sw6ev/).

#### *a. Westerhausen et al. (2004, 2006) sample*

The original publication only published mean corpus callosum area data of a subsample: (a) Westerhausen et al. (2004) included 67 consistent left- and right-handed individuals for which both DTI and morphological corpus callosum measures were available; (b) Westerhausen et al. (2006) included 22 additional participants, which did not show consistent hand preference. The present analysis extends these previous two reports by also including participants for which no DTI data was available, so that the total N = 147.

Hand preference was assessed using a modified German version of the Edinburgh Handedness Inventory (EHI) including 9 items answered on a 5-point scale. Answers were scored from −2 for "always left" to +2 for "always right," yielding a total score ranging from -18 to 18.

For meta-analysis 1, a total score of 0 was used as cut-off to split the sample into dominant left- (dLH) and right-handed (dRH) individuals. Considering meta analyses 2 to 4, a total score of -14.4 or smaller and 14.4 or larger (representing 80% of the maximum absolute score) was applied to define consistent left- (cLH) and right-handed (cRH) individuals, respectively. This cut-off follows the strategy reported by Habib et al. (1991), one of the few studies which reported significant group differences. Individuals with a total score between these values were considered mixed-handed (MH). The group of non-consistent right-handers (NcRH) was formed by joining MH and cLH participants.

The resulting number of subjects (N), mean callosal areas ($cm^2$), and standard deviation (sd) for the groups were:

| Group | N | mean | sd |
|---|---|---|---|
| dRH | 63 | 6.167 | 1.039 |
| dLH | 84 | 5.841 | 0.817 |
| cRH | 51 | 6.118 | 0.970 |
| MH | 46 | 5.907 | 1.044 |
| cLH | 50 | 5.909 | 0.764 |
| NcRH | 96 | 5.908 | 0.904 |
| Total | 147 | 5.981 | 0.930 |

The correlation of the hand preference score with total callosal area was $r = .166$, $p = .045$. Using the absolute score the correlation was $r = .075$, $p = .368$.

*b. Van der Haegen et al. (2011) sample*

The available raw data includes a subsample of N=98 participants from the Van der Haegen et al. (2011) study for which corpus callosum data was available. The corpus callosum is assessed, as midsagittal surface area measures have not been published previously but were made available via author contact. Hand preference for this sample was assessed using the EHI questionnaire and the ariable coding hand prereference ranged range from -3 (all activities done with the left hand) to 3 (all activities done with the right hand).

Regarding meta-analysis 1, a total score of 0 was used to form a dLH and a dRH group. For meta-analyses 2 to 4, a total score of -2.4 or smaller and 2.4 or larger (representing 80% of the maximum absolute test score) was applied to cLH and cRH individuals, respectively. A total score between these values led to a classification of the indivuals as mixed-handed (MH). MH and cLH participants were joined to form the NcRH group.

Applying the above criteria the following number of subjects (N), mean callosal areas ($cm^2$), and standard deviation (sd) were obtained:

| Group | N | mean | sd |
|---|---|---|---|
| dRH | 30 | 4.888 | 0.738 |
| dLH | 68 | 4.923 | 0.653 |
| cRH | 24 | 4.868 | 0.581 |
| MH | 32 | 5.044 | 0.764 |
| cLH | 42 | 4.814 | 0.710 |
| NcRH | 74 | 4.944 | 0.667 |
| Total | 98 | 4.912 | 0.676 |

Hand preference score correlated with callosal area at $r = .003$, $p = .975$. The correlation with absolute $r = .020$, $p = .846$.

*c. Häberling et al., (2011, 2012) sample*

The first author provided the mean values for the two publications (Haberling et al., 2011, 2012) by personal communication on February 12, 2021.

Handedness was assessed with the Annett inventory (range -100% to 100%). A cut-off of 0 points was used to form the dRH and dLH samples and an index of larger 80 and smaller -80% was used to define cRH and cLH groups, respectively. MH included all participants between these values.

The following number of subjects (N), mean callosal areas ($cm^2$), and standard deviation (sd) was obtained for the 2011 publication:

| Group | N | mean | sd |
|---|---|---|---|
| dRH | 35 | 6.419 | 0.814 |
| dLH | 25 | 6.440 | 0.985 |
| cRH | 21 | 6.362 | 0.692 |
| MH | 24 | 6.387 | 0.807 |
| cLH | 15 | 6.585 | 1.218 |

The values for the 2012 publication were:

| Group | N | mean | sd |
|---|---|---|---|
| dRH | 53 | 6.699 | 0.874 |
| dLH | 17 | 6.833 | 0.838 |
| cRH | 41 | 6.594 | 0.719 |
| MH | 21 | 6.855 | 1.074 |
| cLH | 8 | 7.110 | 0.859 |

For the 2011 study, hand preference score correlated with callosal area at $r = -.055$, $p = .679$. For the 2012 study, it was $r = -.141$, $p = .246$

*d. McDowell et al. (2016) sample*

The obtained raw data includes a sample of N = 161 datasets from McDowell et al. (2016), therefore is smaller than the sample reported in the original publication (N = 164). Handedness was assessed using the 5-item Bryden questionnaire, and the laterality score reached from -1 (consistent left) to 1 (consistent right hander). The original publication compared all consistent individuals (i.e. scoring 1 or -1) with all others individuals.

Re-analysing the data for the meta-analyses, a total score of 0 was used as a cut-off value separating the dLH and dRH groups. A total score of -0.80 or smaller and 0.80 or larger (i.e. 80% of the maximum test score) was used to define cLH and cRH handers, respectively. A score between these values led to a classification of individuals as mixed-handed (MH). MH and cLH groups taken together formed the NcRH group.

Of note, the dependent variable was total corpus callosum volume, obtained using Freesurfer segmentation so that mean corpus callosum is expressed as volume ($cm^3$):

| Group | N | mean | sd |
|---|---|---|---|
| dRH | 143 | 3150.00 | 323.93 |
| dLH | 18 | 3149.08 | 441.76 |
| cRH | 119 | 3141.02 | 436.56 |
| MH | 36 | 3172.64 | 452.76 |
| cLH | 9 | 3123.89 | 220.15 |

| | | | |
|---|---|---|---|
| NcRH | 45 | 3162.89 | 415.05 |
| Total | 164 | 3147.02 | 429.61 |

Hand preference score and callosal volume correlated at $r = -.008$, $p = .915$. The correlation with the absolute value of the hand preference score was $r = -.018$, $p = .816$.

#### *e. Labache et al. (2020) sample*

The sample includes N=287 datasets as published in Labache et al. (2020), whereby the corpus callosum data has previously not been analyzed regarding hand preference. Hand preference was assessed using the EHI and the total score ranged from LI = -100 (all activities done with the left hand) to 100 (all activities done with the right hand).

For meta-analysis 1 a total score of 0 was used as a cut-off value separating the dLH and dRH groups, whereby 8 participants with the total score of 0 were excluded. For meta-analyses 2 to 4, a total score of -80 or smaller and 80 or larger (i.e. 80% of the maximum absolute test score) was used to define cLH and c>RH individuals, respectively. A score between these values led to a classification of the individuals as mixed-handed (MH). MH and cLH groups taken together formed the NcRH group.

Following the above classification criteria the following number of subjects (N), mean corpus callosum volume ($cm^3$), and standard deviation (sd) were obtained:

| **Group** | **N** | **Mean** | **sd** |
|---|---|---|---|
| dRH | 148 | 5.346 | 0.828 |
| dLH | 131 | 5.313 | 0.857 |
| cRH | 118 | 5.362 | 0.828 |
| MH | 105 | 5.188 | 1.006 |
| cLH | 64 | 5.484 | 0.857 |
| NcRH | 169 | 5.300 | 0.909 |
| Total | 287 | 5.325 | 0.844 |

Correlating the LI with total corpus callosum volume yielded an $r = -.022$, $p = .706$. Considering absolute LI values, the correlation was $r = .200$, $p = .001$.

#### *f. Human Connectome Project (Van Essen et al., 2013)*

The total sample includes N = 1113 datasets, which are all participants published in the 1200 Subjects Data Release of the HCP consortium (Van Essen et al., 2013) for which MRI data is available for download in tabulated form. The corpus callosum data represent data-extraction using Freesurfer software (Glasser et al., 2013). Hand preference was assessed using the EHI, and the LQ ranged from -100 (all activities done with the left hand) to 100 (all activities done with the right hand).

Mean and standard deviation of corpus callosum size was determined as follows: an LQ of 0 was used as a cut-off value for separating the dLH and dRH groups (note here 4 participants were excluded as the LQ was 0). An LQ of -80 or smaller and 80 or larger (i.e., 80% of the maximum test score) was used to define cLH and cRH handers. A score between these values led to the classification of the individuals as mixed-handed (MH). The NcRH group was formed by combining MH and cLH groups.

The following number of subjects (N), mean corpus callosum volume ($cm^3$), and standard deviation (sd) were obtained:

| Group | N | Mean | sd |
|---|---|---|---|
| dRH | 1008 | 3327.25 | 479.96 |
| dLH | 101 | 3269.05 | 460.67 |
| cRH | 615 | 3320.97 | 487.35 |
| MH | 470 | 3333.02 | 467.85 |
| cLH | 28 | 3168.14 | 422.62 |
| NcRH | 498 | 3323.75 | 466.59 |
| Total | 1113 | 3322.21 | 477.96 |

The correlation of the LQ with total corpus callosum volume was $r = .034$, $p = .261$; and for absolute LQ the correlation was $r = -.011$, $p = .707$.

### 4. Transfer heuristics between different schemas of callosal subdivision (Figure S1)

| Subdivision | Comparison with reference | Transfer heuristic |
|---|---|---|
| PCA-based segmentation of callosal thickness measures, as used by Denenberg et al. (1991) and Cowell et al. (2018) | 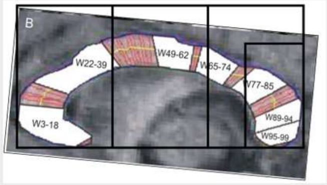  | • W3-18 + W22-39 approximates anterior third<br>• W49-62 approx. middle third<br>• W65-74 approx. Isthmus<br>• W77-85 + W89-94 + W89-94 approx. splenium |
| Radial segmentation used by Habib et al. (1991) | 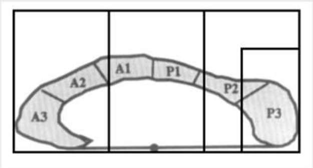  | • A3 + A2 approximates anterior third<br>• A1+P1 approx. middle third<br>• P2 approx. isthmus<br>• P3 approx. splenium |
| Freesurfer segmentation used by McDowell et al. (2016) | 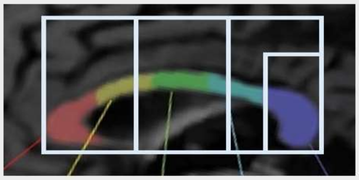 | • Anterior (red) + Midanterior (yellow) approximates anterior third<br>• Central (green) approx mid third<br>• Midposterior (light blue) approx. Isthmus<br>• Posterior (blue) approx Splenium |

*Fig. S1*. Comparison of different subdivision schemas with the Jäncke et al. (1997) subdivision scheme used as reference frame (black and white overlays, see also Fig. 2 main document). Top row shows the PCA based method (Cowell & Gurd, 2018; Denenberg et al., 1991), whereby Fig. 1B from Cowell and Gurd (2018) was used as background. Middle row uses Fig. 1 of Habib et al. (1991) for comparison. Bottom panel, Freesurfer segmentation used by McDowell et al. (2016) and the HCP project (Van Essen et al., 2013), Fig. from Goldman et al., (2017) was used for the comparison. Subregional measures from the original studies were transferred to the reference scheme using the derived heuristics (last column).

### 5. Risk-of-bias analysis within studies

The nature of the topic of the present meta-analysis reduces the risk-of-bias assessment to the "Bias in selection of the reported results" criterion as all other potential criteria are conceptualized for evaluating randomized clinical intervention studies (see Higgins, Savović, Page, Elbers, & Sterne, 2021). All included studies and datasets were scanned for selective reporting, that is, whether potentially available corpus-callosum data was also reported in the results section (evaluator R.W.). This was evaluated separately concerning total and subsection data. The results are presented in the table below.

*Table S6.* Risk-of-bias assessment within studies concerning the data of the total corpus callosum (cc) and the subsection measures

| # | Study | Total cc | Subsection | comment |
|---|---|---|---|---|
| 1 | Clarke and Zaidel (1994) | Low risk of bias | Low risk of bias | All data reported |
| 2 | Cowell and Gurd (2018) | Low risk of bias | Low risk of bias | All data reported |
| 3 | Denenberg et al. (1991) | Low risk of bias | Low risk of bias | All data reported |
| 4 | Haberling et al. (2012) | Low risk of bias | not analysed | All data available after author contact |
| 5 | Haberling et al. (2011) | Low risk of bias | not analysed | All data available after author contact |
| 6 | Habib et al. (1991) | Low risk of bias | Low risk of bias | All data reported |
| 7 | HCP 2017 | Low risk of bias | Low risk of bias | All data available |
| 8 | Hines et al. (1992) | High risk of bias | High risk of bias | Data only reported for sign. comparisons |
| 9 | Jancke et al. (1997) | Low risk of bias | Low risk of bias | All data reported |
| 10 | Kertesz et al. (1987) | Low risk of bias | not analysed | All data reported |
| 11 | Labache et al. (2020) | Low risk of bias | Low risk of bias | All data available |
| 12 | Luders et al. (2003) | Low risk of bias | Low risk of bias | All data reported |
| 13 | Martens et al. (2013) | Low risk of bias | High risk of bias | Selective reporting of sign. subsection data |
| 14 | McDowell et al. (2016) | Low risk of bias | Low risk of bias | All data available |
| 15 | Moffat et al. (1998) | Low risk of bias | High risk of bias | Data only reported for sign. comparisons |
| 16 | Morton and Rafto (2006) | Low risk of bias | not analysed | All data reported |
| 17 | Nasrallah et al. (1986) | High risk of bias | not analysed | Only data of male sample reported |
| 18 | Ozdikici (2020) | Low risk of bias | not analysed | All data reported |

| | | | | |
|---|---|---|---|---|
| 19 | Steinmetz et al. (1992) | Low risk of bias | Some concerns | Data of anterior regions not reported for theoretical reasons |
| 20 | Tuncer et al. (2005) | Low risk of bias | High risk of bias | Subregion not reported although analysed |
| 21 | Van der Haegen et al. (2011) | Low risk of bias | Low risk of bias | All data available |
| 22 | Westerhausen et al. (2004) | Low risk of bias | Low risk of bias | All data available |
| 23 | Witelson (1985) | Low risk of bias | Low risk of bias | All data reported |
| 24 | Witelson (1989) | Low risk of bias | Low risk of bias | All data reported |
| 25 | Witelson and Goldsmith (1991) | Low risk of bias | Some concerns | Data only of isthmus reported for theoretical reasons |

### 6. Supplement Figure S2: Funnel plots of main analyses

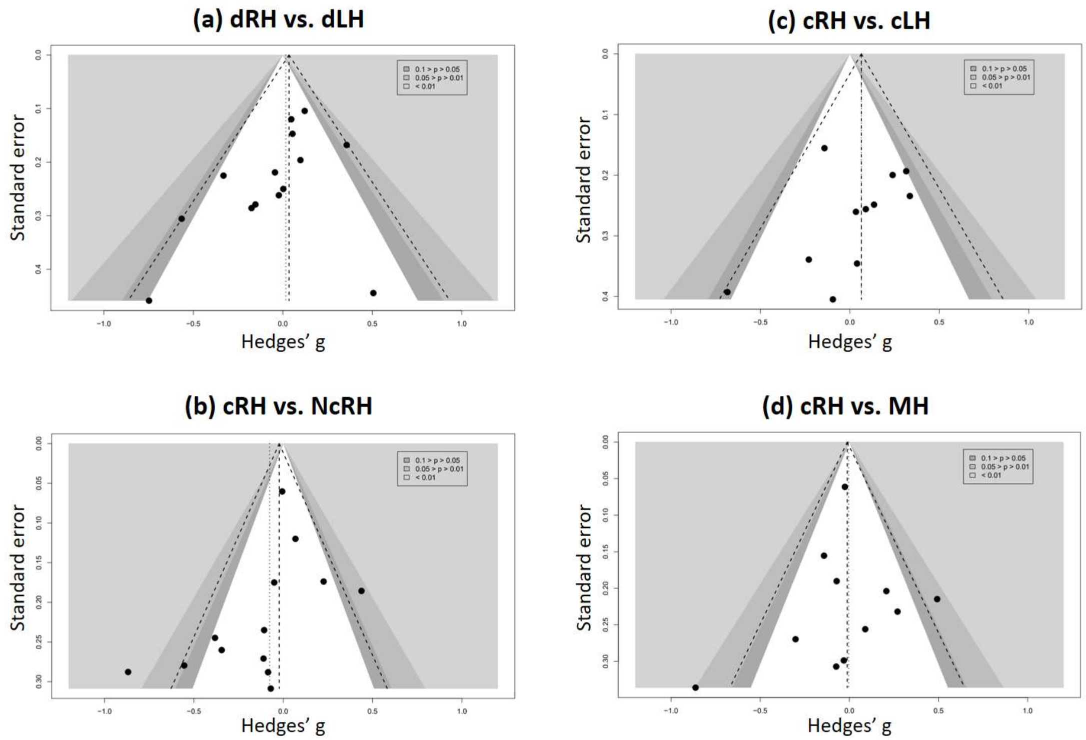

*Fig. S2*. Funnel plots of the four meta-analyses relating hand preference to total corpus callosum size (for details see text). Note that the shaded areas reflect the level of significance as indicated in the legend of each plot

### 7. Supplement Figure S3: moderator analysis by sex

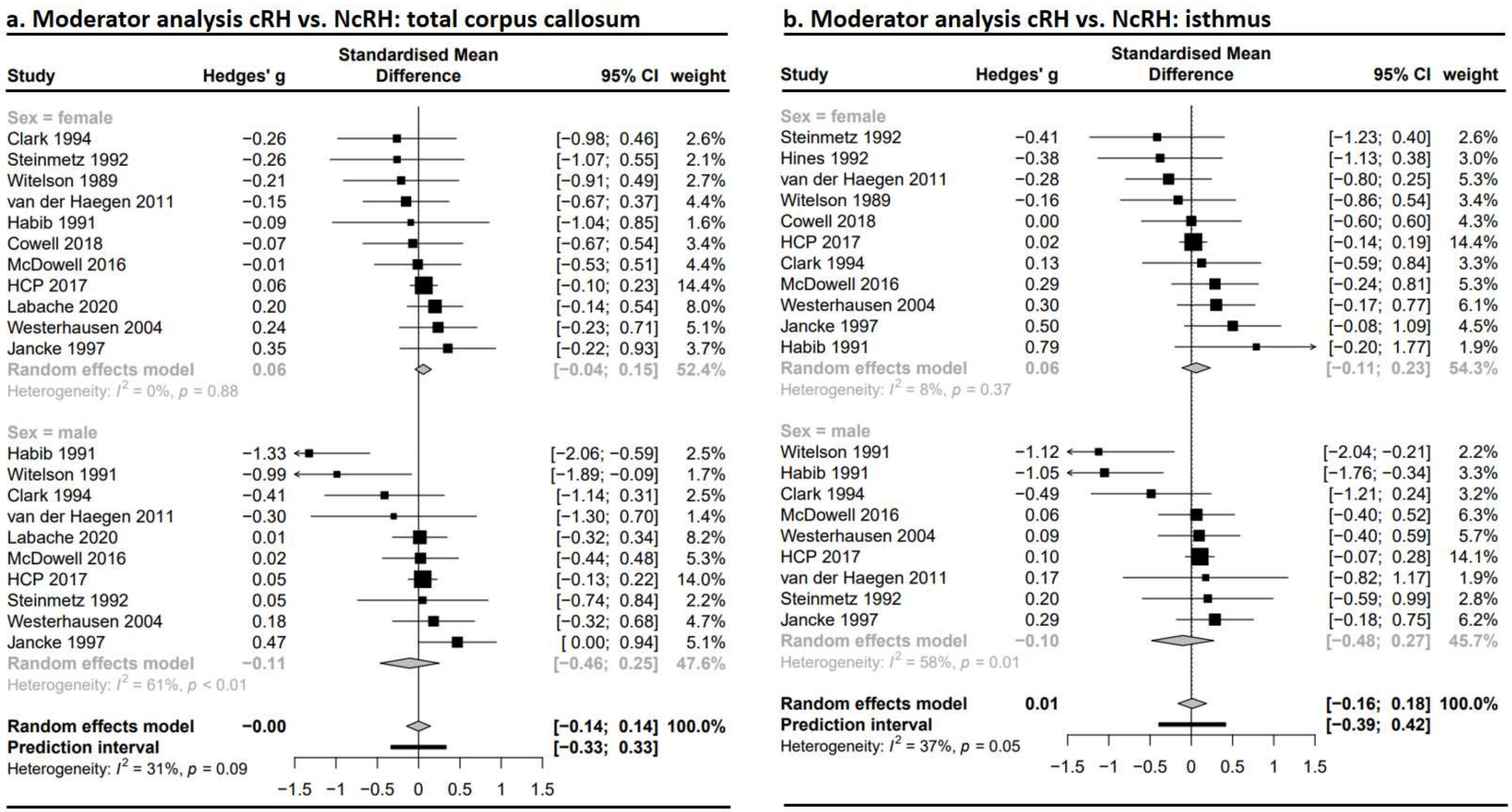

*Fig. S3*. Forest plots of the moderator analysis by sex. The results for the total corpus callosum are presented in (a) and for the isthmus in (b). More details regarding the main analysis can be found in the main text.

### 8. Supplement Figure S4: overview meta-analysis by subsection for cRH vs. cLH and cRH vs. MH comparisons.

| | | Anterior third | | Middle third | | Isthmus | | Splenium | |
|---|---|---|---|---|---|---|---|---|---|
| Comarison | Study | Hedges' g | se(g) | Hedges' g | se(g) | Hedges' g | se(g) | Hedges' g | se(g) |
| **dRH vs. dLH** | Denenberg 1991 | -0.01 | 0.20 | 0.06 | 0.20 | 0.30 | 0.20 | 0.02 | 0.20 |
| | HCP 2017 | 0.16 | 0.10 | 0.08 | 0.10 | 0.11 | 0.10 | 0.02 | 0.10 |
| | Martens 2013 | n.a. | n.a. | -0.74 | 0.46 | n.a. | n.a. | n.a. | n.a. |
| | Moffat 1998 | n.a. | n.a. | -0.51 | 0.30 | -0.97 | 0.32 | n.a. | n.a. |
| | McDowell 2016 | 0.02 | 0.25 | -0.05 | 0.25 | 0.19 | 0.25 | -0.12 | 0.25 |
| | vdHaegen 2011 | -0.04 | 0.22 | 0.02 | 0.22 | -0.16 | 0.22 | -0.04 | 0.22 |
| | Westerhausen 2004 | 0.40 | 0.17 | 0.28 | 0.17 | 0.26 | 0.17 | 0.30 | 0.17 |
| | Average [CI] effect | n.d. | n.d. | n.d. | n.d. | n.d. | n.d | n.d. | n.d. |
| **cRH vs. cLH** | HCP 2017 | 0.31 | 0.19 | 0.09 | 0.19 | 0.13 | 0.19 | 0.41 | 0.19 |
| | Jäncke 1997 | 0.28 | 0.23 | 0.43 | 0.24 | 0.33 | 0.23 | 0.13 | 0.23 |
| | Lüders 2003 | 0.10 | 0.26 | 0.00 | 0.26 | -0.33 | 0.26 | 0.07 | 0.26 |
| | McDowell 2016 | 0.10 | 0.35 | 0.06 | 0.35 | 0.24 | 0.35 | -0.21 | 0.35 |
| | Steinmetz 1992 | n.a. | n.a. | 0.07 | 0.40 | 0.00 | 0.40 | -0.09 | 0.40 |
| | vdHaegen 2011 | -0.04 | 0.26 | -0.06 | 0.26 | -0.17 | 0.26 | -0.05 | 0.26 |
| | Westerhausen 2004 | 0.31 | 0.20 | 0.28 | 0.20 | 0.22 | 0.20 | 0.04 | 0.20 |
| | Average [CI] effect | n.d. | n.d. | n.d. | n.d. | n.d. | n.d. | n.d. | n.d. |
| **cRH vs. MH** | HCP 2017 | -0.03 | 0.06 | -0.04 | 0.06 | 0.03 | 0.06 | -0.03 | 0.06 |
| | Jäncke 1997 | 0.26 | 0.21 | 0.63 | 0.22 | 0.37 | 0.21 | 0.48 | 0.21 |
| | McDowell 2016 | -0.08 | 0.19 | -0.08 | 0.19 | 0.09 | 0.19 | -0.09 | 0.19 |
| | Steinmetz 1992 | n.a. | n.a. | 0.08 | 0.31 | -0.19 | 0.31 | -0.06 | 0.31 |
| | vdHaegen 2011 | -0.26 | 0.27 | -0.18 | 0.27 | -0.17 | 0.27 | -0.46 | 0.27 |
| | Westerhausen 2004 | 0.13 | 0.20 | 0.20 | 0.20 | 0.20 | 0.20 | 0.29 | 0.20 |
| | Witelson 1985 | n.a. | n.a. | n.a. | n.a. | n.a. | n.a. | -0.41 | 0.33 |
| | Average [CI] effect | n.d. | n.d. | n.d. | n.d. | n.d. | n.d. | **0.00** | **[-0.28; 0.28]** |

| **Hedges' g** |
|---|
| >0.80 |
| >0.50 |
| >0.20 |
| >0 |
| 0 |
| <0 |
| <-0.20 |
| <-0.50 |
| <-0.80 |

*Fig. S4.* Presents the effect size (*g*) and standard error of the effect size (*se(g)*) for each study included in the comparison of dRH and dLH (top), cRH and cLH (middle), and cRH and MH (bottom), respectively. Negative values indicate the subsection to be larger in the non-right-handed group (dLH, cLH, MH), positive values indicate the dRH/cRH group to have a larger callosal subsection. A meta-analytic average was only calculated were a sufficient amount of studies was available (as determined by power analysis, see main text), and otherwise not determined (n.d.). The provided mean effect is determined within a random-effects model. The values in brackets are the 95% confidence interval. Color coding was based on the Cohen's effect-size heuristics (Cohen, 1992) as indicated in the figure legend. Note, for some studies data was not available (n.a.) for some of the subsection.

### 9. Supplementary analysis: meta-regression differential sex distribution

As sex is associated with differences in the absolute size of the corpus callosum (Bishop & Wahlsten, 1997; Smith, 2005), a difference in the distribution of males and females in the compared handedness groups might potentially confound the comparison of the effect sizes.

To test for this possibility, we conducted a meta-regression analysis for the comparisons dRH vs. dLH and cRH vs. NcRH using the difference in the proportion of females as covariate. That is, a variable *deltaF* was determined as the percentage of female in the right-handed group minus the percentage of females in the non-right handed group (see Tables S1 and S2 for the data basis). Thus, positive values indicated a larger proportion of females in the right-handed sample and negative values a larger proportion of females in the non-right handed sample. Given that males can be expected to have the larger absolute corpus callosum, a larger proportion of females should affect the mean corpus callosum size in the respective group negatively. Consequently, a negative association of deltaF with the effect size would be indicative of the suspected confounding effect. Note, the sex distribution by sample was not available for all studies included in the meta-analysis so that the number of the studies included in the meta-regression analysis is reduced compared with the respective meta-analysis. The analysis was conducted using the *metareg* function of the *metafor* R library (Viechtbauer, 2020).

The meta-regression coefficient was $\beta = 0.218$ with a $t = 0.11$, $p = 0.91$ for the dRH vs. dLH analysis and including $k = 11$ studies. The coefficient was $\beta = 0.112$ with $t = 0.10$, $p = 0.92$ for the cRH vs. NcRH effect sizes ($k = 11$ studies). In both cases the percentage explained variance by deltaF was below 0.001%.

### 10. Supplementary analysis: absolute LQ

Spearman correlation ($r_{sp}$) for the association of |LQ| and total corpus callosum area or volume. Rank correlation were considered appropriate as the distribution of |LQ| is non-normal. The data from Habib et al. (1991) was reported in Table 2 (p. 49) of this publication. All other correlation were calculated from the raw data available (see Supplement Section 3).

| Study | N | $r_{sp}$ | *p* |
|---|---|---|---|
| Habib et al. (1991) | 53 | 0.297 | 0.03 |
| HCP data | 1113 | -0.013 | 0.67 |
| Labache et al. (2020) | 287 | 0.099 | 0.09 |
| McDowell et al. (2016) | 164 | -0.016 | 0.84 |
| Van der Haegen et al. (2011) | 98 | -0.154 | 0.15 |
| Westerhausen et al. (2004) | 147 | 0.073 | 0.38 |

### References

Bishop, K. M., & Wahlsten, D. (1997). Sex differences in the human corpus callosum: Myth or reality? *Neuroscience and Biobehavioural Reviews, 21*(5), 581-601.

Burke, H. L., & Yeo, R. A. (1994). Systematic variations in callosal morphology: The effects of age, gender, hand preference, and anatomic asymmetry. *Neuropsychology, 8*(4), 563-571.

Cherbuin, N., Luders, E., Chou, Y. Y., Thompson, P. M., Toga, A. W., & Anstey, K. J. (2013). Right, left, and center: How does cerebral asymmetry mix with callosal connectivity? *Human Brain Mapping, 34*(7), 1728-1736. doi:10.1002/hbm.22022

Clarke, J. M., Lufkin, R. B., & Zaidel, F. (1993). Corpus-callosum morphometry and dichotic-listening perfromance - individual diiderences in functional interhemispheric inhibition. *Neuropsychologia, 31*(6), 547-557. doi:10.1016/0028-3932(93)90051-z

Clarke, J. M., & Zaidel, E. (1994). Anatomical-behavioral relationships: Corpus callosum morphometry and hemispheric specialization. *Behavioural Brain Research, 64*(1-2), 185-202. doi:10.1016/0166-4328(94)90131-7

Cohen, J. (1992). A power primer. *Psychological Bulletin, 112*(1), 155.

Cowell, P. E., & Gurd, J. (2018). Handedness and the corpus callosum: A review and further analyses of discordant twins. *Neuroscience, 388*, 57-68. doi:10.1016/j.neuroscience.2018.06.017

Cowell, P. E., Kertesz, A., & Denenberg, V. H. (1993). Multiple dimensions of handedness and the human corpus callosum. *Neurology, 43*(11), 2353-2357. doi:10.1212/wnl.43.11.2353

Denenberg, V. H., Kertesz, A., & Cowell, P. E. (1991). A factor analysis of the human's corpus callosum. *Brain Research, 548*(1-2), 126-132. doi:10.1016/0006-8993(91)91113-f

Glasser, M. F., Sotiropoulos, S. N., Wilson, J. A., Coalson, T. S., Fischl, B., Andersson, J. L., . . . HCP-Consortium. (2013). The minimal preprocessing pipelines for the Human Connectome Project. *Neuroimage, 80*, 105-124.

Goldman, J. G., Bledsoe, I. O., Merkitch, D., Dinh, V., Bernard, B., & Stebbins, G. T. (2017). Corpus callosal atrophy and associations with cognitive impairment in Parkinson disease. *Neurology, 88*(13), 1265-1272.

Gurd, J. M., Cowell, P. E., Lux, S., Rezai, R., Cherkas, L., & Ebers, G. C. (2013). fMRI and corpus callosum relationships in monozygotic twins discordant for handedness. *Brain Structure & Function, 218*(2), 491-509. doi:10.1007/s00429-012-0410-9

Haberling, I. S., Badzakova-Trajkov, G., & Corballis, M. C. (2011). Callosal tracts and patterns of hemispheric dominance: A combined fMRI and DTI study. *Neuroimage, 54*(2), 779-786. doi:10.1016/j.neuroimage.2010.09.072

Haberling, I. S., Badzakova-Trajkov, G., & Corballis, M. C. (2012). The corpus callosum in monozygotic twins concordant and discordant for handedness and language dominance. *Journal of Cognitive Neuroscience, 24*(10), 1971-1982. Retrieved from <Go to ISI>://WOS:000308422200001

Habib, M., Gayraud, D., Oliva, A., Regis, J., Salamon, G., & Khalil, R. (1991). Effects of handedness and sex on the morphology of the corpus callosum: A study with brain magnetic resonance imaging. *Brain and Cognition, 16*(1), 41-61.

Higgins, J., Savović, J., Page, M., Elbers, R., & Sterne, J. (2021). Chapter 8: Assessing risk of bias in a randomized trial. In J. Higgins, J. Thomas, J. Chandler, M. Cumpston, T. Li, M. Page, & V. Welch (Eds.), *Cochrane Handbook for Systematic Reviews of Interventions version 6.2 (updated February 2021).* : Cochrane.

Hines, M., Chiu, L., McAdams, L. A., Bentler, P. M., & Lipcamon, J. (1992). Cognition and the corpus callosum: Verbal fluency, visuospatial ability, and language lateralization related to midsagittal surface areas of callosal subregions. *Behavioral Neuroscience, 106*(1), 3.

Hopper, K. D., Patel, S., Cann, T. S., Wilcox, T., & Schaeffer, J. M. (1994). The relationship of age, gender, handedness, and sidedness to the size of the corpus callosum. *Academic Radiology, 1*(3), 243-248. doi:10.1016/s1076-6332(05)80723-8

Jancke, L., Staiger, J. F., Schlaug, G., Huang, Y. X., & Steinmetz, H. (1997). The relationship between corpus callosum size and forebrain volume. *Cerebral Cortex, 7*(1), 48-56. doi:10.1093/cercor/7.1.48

Josse, G., Seghier, M. L., Kherif, F., & Price, C. J. (2008). Explaining function with anatomy: Language lateralization and corpus callosum size. *Journal of Neuroscience, 28*(52), 14132-14139. doi:10.1523/jneurosci.4383-08.2008

Kertesz, A., Polk, M., Howell, J., & Black, S. E. (1987). Cerebral dominance, sex, and callosal size in MRI. *Neurology, 37*(8), 1385-1388. doi:10.1212/wnl.37.8.1385

Labache, L., Mazoyer, B., Joliot, M., Crivello, F., Hesling, I., & Tzourio-Mazoyer, N. (2020). Typical and atypical language brain organization based on intrinsic connectivity and multitask functional asymmetries. *Elife, 9*. doi:10.7554/eLife.58722

Luders, E., Cherbuin, N., Thompson, P. M., Gutman, B., Anstey, K. J., Sachdev, P., & Toga, A. W. (2010). When more is less: Associations between corpus callosum size and handedness lateralization. *Neuroimage, 52*(1), 43-49. doi:10.1016/j.neuroimage.2010.04.016

Luders, E., Rex, D. E., Narr, K. L., Woods, R. P., Jancke, L., Thompson, P. M., . . . Toga, A. W. (2003). Relationships between sulcal asymmetries and corpus callosum size: Gender and handedness effects. *Cerebral Cortex, 13*(10), 1084-1093. doi:10.1093/cercor/13.10.1084

Martens, M. A., Wilson, S. J., Chen, J., Wood, A. G., & Reutens, D. C. (2013). Handedness and corpus callosal morphology in Williams syndrome. *Development and Psychopathology, 25*(1), 253-260. doi:10.1017/s0954579412001009

McDowell, A., Felton, A., Vazquez, D., & Chiarello, C. (2016). Neurostructural correlates of consistent and weak handedness. *Laterality, 21*(4-6), 348-370. doi:10.1080/1357650x.2015.1096939

Mitchell, T. N., Free, S. L., Merschhemke, M., Lemieux, L., Sisodiya, S. M., & Shorvon, S. D. (2003). Reliable callosal measurement: Population normative data confirm sex-related differences. *American Journal of Neuroradiology, 24*(3), 410-418.

Moffat, S. D., Hampson, E., & Lee, D. H. (1998). Morphology of the planum temporale and corpus callosum in left handers with evidence of left and right hemisphere speech representation. *Brain, 121*, 2369-2379. doi:10.1093/brain/121.12.2369

Morton, B. E., & Rafto, S. E. (2006). Corpus callosum size is linked to dichotic deafness and hemisphericity, not sex or handedness. *Brain and Cognition, 62*(1), 1-8. doi:10.1016/j.bandc.2006.03.001

Nasrallah, H. A., Andreasen, N. C., Coffman, J. A., Olson, S. C., Dunn, V. D., Ehrhardt, J. C., & Chapman, S. M. (1986). A controlled magnetic resonance imaging study of corpus callosum thickness in schizophrenia. *Biological Psychiatry, 21*(3), 274-282. doi:10.1016/0006-3223(86)90048-x

O'Kusky, J., Strauss, E., Kosaka, B., Wada, J., Li, D., Druhan, M., & Petrie, J. (1988). The corpus callosum is larger with right-hemisphere cerebral speech dominance. *Annals of Neurology: Official Journal of the American Neurological Association and the Child Neurology Society, 24*(3), 379-383.

Ozdikici, M. (2020). Measurment of midsagittal corpus callosum area with the modified Cavalieri method in healthy right- and left-handed Turkish adults. *Malang Neurology Journal, 6*(1), 24-27.

Preuss, U. W., Meisenzahl, E. M., Frodl, T., Zetzsche, T., Holder, J., Leinsinger, G., . . . Möller, H. J. (2002). Handedness and corpus callosum morphology. *Psychiatry Research, 116*(1-2), 33-42. doi:10.1016/s0925-4927(02)00064-1

Reinarz, S. J., Coffman, C. E., Smoker, W. R., & Godersky, J. C. (1988). MR imaging of the corpus callosum: Normal and pathologic findings and correlation with CT. *AJR Am J Roentgenol, 151*(4), 791-798. doi:10.2214/ajr.151.4.791

Robichon, F., & Habib, M. (1998). Abnormal callosal morphology in male adult dyslexics: Relationships to handedness and phonological abilities. *Brain and Language, 62*(1), 127-146. doi:10.1006/brln.1997.1891

Smith, R. J. (2005). Relative size versus controlling for size. *Current Anthropology, 46*(2), 249-273.

Steinmetz, H., Jäncke, L., Kleinschmidt, A., Schlaug, G., Volkmann, J., & Huang, Y. (1992). Sex but no hand difference in the isthmus of the corpus callosum. *Neurology, 42*(4), 749-752. doi:10.1212/wnl.42.4.749

Steinmetz, H., Staiger, J. F., Schlaug, G., Huang, Y., & Jäncke, L. (1995). Corpus callosum and brain volume in women and men. *Neuroreport, 6*(7), 1002-1004. doi:10.1097/00001756-199505090-00013

Tuncer, M. C., Hatipoglu, E. S., & Ozates, M. (2005). Sexual dimorphism and handedness in the human corpus callosum based on magnetic resonance imaging. *Surgical and Radiological Anatomy, 27*(3), 254-259. doi:10.1007/s00276-004-0308-1

Van der Haegen, L., Cai, Q., Seurinck, R., & Brysbaert, M. (2011). Further fMRI validation of the visual half field technique as an indicator of language laterality: A large-group analysis. *Neuropsychologia, 49*(10), 2879-2888.

Van Essen, D. C., Smith, S. M., Barch, D. M., Behrens, T. E., Yacoub, E., Ugurbil, K., & HCP-Consortium. (2013). The WU-Minn human connectome project: An overview. *Neuroimage, 80*, 62-79.

Viechtbauer, W. (2020). metafor: Meta-Analysis Package for R. Retrieved from https://CRAN.R-project.org/package=metafor

Welcome, S. E., Chiarello, C., Towler, S., Halderman, L. K., Otto, R., & Leonard, C. M. (2009). Behavioral correlates of corpus callosum size: Anatomical/behavioral relationships vary across sex/handedness groups. *Neuropsychologia, 47*(12), 2427-2435. doi:10.1016/j.neuropsychologia.2009.04.008

Westerhausen, R., Kreuder, F., Dos Santos Sequeira, S. D., Walter, C., Woerner, W., Wittling, R. A., . . . Wittling, W. (2004). Effects of handedness and gender on macro- and microstructure of the corpus callosum and its subregions: A combined high-resolution and diffusion-tensor MRI study. *Cognitive Brain Research, 21*(3), 418-426. doi:10.1016/j.cogbrainres.2004.07.002

Westerhausen, R., Kreuder, F., Sequeira, S. D. S., Walter, C., Woerner, W., Wittling, R. A., . . . Wittling, W. (2006). The association of macro-and microstructure of the corpus callosum and language lateralisation. *Brain and Language, 97*(1), 80-90.

Witelson, S. F. (1985). The brain connection: The corpus callosum is larger in left-handers. *Science, 229*(4714), 665-668. doi:10.1126/science.4023705

Witelson, S. F. (1989). Hand and sex differences in the isthmus and genu of the human corpus callosum. A postmortem morphological study. *Brain, 112 ( Pt 3)*, 799-835. doi:10.1093/brain/112.3.799

Witelson, S. F., & Goldsmith, C. H. (1991). The relationship of hand preference to anatomy of the corpus callosum in men. *Brain Research, 545*(1-2), 175-182. doi:10.1016/0006-8993(91)91284-8
